# Upregulated SEMA3C in astrocytes contributes to Rett Syndrome phenotypes

**DOI:** 10.1101/2025.10.10.681027

**Authors:** Krissy A. Lyon, Adrien Paumier, Ananya Kandikonda, Andrea Melendez, Nicola J. Allen

## Abstract

Astrocytes support neuronal function during development through secreted proteins, yet how astrocyte-secreted cues are altered in disease states and contribute to neurodevelopmental disorders remains poorly defined. Rett syndrome (RTT) is a regressive neurodevelopmental disorder characterized by motor, sensory, and cognitive impairments. Here, we identify the class 3 semaphorin SEMA3C as an astrocyte-secreted protein that contributes to RTT pathology. We show that *Sema3c* expression is elevated in astrocytes *in vivo* in RTT model mice and find that SEMA3C is sufficient to inhibit cortical neuron dendrite outgrowth. Normalization of astrocyte SEMA3C levels in female RTT model mice rescues dendritic arborization deficits, restores synaptic activity, and improves visual acuity and motor behavior. Mechanistically, both SEMA3C and RTT astrocyte conditioned media inhibit dendrite outgrowth through PLXND1-dependent signaling. Together, these findings identify astrocyte-secreted SEMA3C as a contributor to RTT pathology and highlight SEMA3C and PLXND1 signaling as potential therapeutic targets in neurodevelopmental disorders.

## Introduction

Rett syndrome (RTT) is an X-linked neurodevelopmental disorder involving loss-of-function of the methyl-CpG-binding protein 2 (*MECP2*) gene resulting in regressive motor, visual, and cognitive deficits^1,2^. Mutations in MECP2, a transcriptional regulator, impact gene expression critical for nervous system development and neuronal connectivity^3–6^. While RTT research has primarily focused on neurons, it is now clear that astrocytes contribute to RTT pathology^7–17^. Co-culture of RTT astrocytes with wildtype neurons reduces dendritic outgrowth, suggesting a non-cell autonomous effect of astrocytes through secreted factors^15,17^. Proteomic analysis of astrocyte secreted factors in RTT identified numerous proteins with aberrant secretion^10,12,18^. Yet, if and how these altered astrocyte secreted factors contribute to RTT pathology is largely unknown.

During development, astrocytes secrete proteins that modulate synapse formation, stabilization, and connectivity^19^. In neurodevelopmental disorders, astrocyte dysfunction contributes to altered neuronal function^12,13,15,18,20–30^. Individuals with RTT show dysfunction across visual, auditory, and somatosensory systems, reflected in altered evoked neuronal responses, oscillations, and functional connectivity^31–40^. Postmortem tissue shows smaller neurons with reduced dendritic arborization in individuals with RTT^-41,42^. RTT mouse models show similar neuronal morphology changes and altered synaptic function^42–51^. Astrocyte-specific restoration of MECP2 in otherwise MECP2-null mice improves dendrite morphology, locomotion, anxiety, and respiratory phenotypes, and prolongs lifespan^16^. While restoration of MECP2 expression is one potential therapeutic approach^52–55^, overexpression of MECP2 can lead to deficits^56^. MECP2 duplication syndrome is a condition associated with intellectual disability and seizures, primarily affecting males. In mouse models, overexpression of MECP2 results in similar outcomes including motor deficits, anxiety, and seizures^57,58^. These challenges of gene dosage sensitivity highlight the importance of pursuing alternative approaches to identify and modulate factors that contribute to deficits in RTT as potential therapeutic targets.

Towards linking specific astrocyte secreted proteins to morphological and functional changes, this study focuses on the impacts of increased astrocyte SEMA3C on RTT deficits. Proteomic analysis of astrocytes from models of RTT and two other neurodevelopmental disorders, Fragile X syndrome and Down syndrome, revealed increased secretion of the class 3 semaphorin, SEMA3C^12^. Class 3 semaphorins (SEMA3A-G) are secreted proteins first characterized for their role in axon guidance^59^, with more recent evidence for roles in dendrite outgrowth, spine formation, and synaptogenesis^60–64^. Moreover, members of the semaphorin family are implicated in astrocyte regulation of neurodevelopment^65,66^. In RTT, transcriptomic and proteomic data reveal dysregulation of semaphorins, and their receptors neuropilins and plexins^67,68^, in human^18,69,70^ and mouse samples^10,12,69,71–74^. Further, acute loss of MECP2 in adult mice alters semaphorin signaling^72^, and SEMA3F dysfunction is associated with altered neuronal connectivity in the olfactory bulb and hippocampus in *Mecp2*-null mice^75,76^. While transcriptomic and proteomic datasets point to altered semaphorin signaling in RTT, few studies to date have explored semaphorin impacts on RTT phenotypes. We investigate this question using a female *Mecp2* heterozygous mouse model. Since *MECP2* resides on the X chromosome, individuals diagnosed with RTT are predominantly female^1^, and random X inactivation produces mosaic *MECP2* expression, with some cells expressing wildtype *MECP2* and others expressing mutant *MECP2.* Female *Mecp2* heterozygous mouse models capture this mosaic cellular environment and allow for cellular changes to be linked to behavioral outcomes^23,49,53,77–86^.

Here, by combining experiments in astrocyte and neuron cell culture, with genetic reduction of astrocyte SEMA3C in RTT model mice, we find that upregulated astrocyte SEMA3C contributes to RTT deficits. Using cell culture, we show that SEMA3C is inhibitory to cortical neuron dendrite outgrowth. To ask if SEMA3C contributes to RTT phenotypes *in vivo*, we normalized astrocyte SEMA3C to wildtype levels in *Mecp2* heterozygous mice, finding this rescues neuronal morphology, synaptic activity, and behavior deficits. We find that both SEMA3C and RTT astrocyte conditioned media inhibit dendrite outgrowth through PLXND1-dependent signaling. Our findings identify astrocytic upregulation of SEMA3C as one mechanism by which astrocytes contribute to RTT pathogenesis and highlight semaphorin-plexin signaling as an important regulator of astrocyte-neuron interactions in neurodevelopment and disease.

## Results

### SEMA3C inhibits cortical neuron dendrite outgrowth

Astrocyte protein secretion of SEMA3C is increased in RTT^12^ (Figure 1A). SEMA3C mediates chemoattraction or repulsion in different cellular and developmental contexts^87–91^. To understand how increased SEMA3C secretion impacts neurons, we first used astrocyte and neuron cell culture to test the effect of SEMA3C on neuron dendrite outgrowth. First, we applied wildtype (WT) postnatal day (P)7 cortical astrocyte conditioned media (ACM) to age-matched cortical neurons with and without recombinant SEMA3C protein (Figures 1B and 1C). Immunostaining for MAP2 to label dendrites after three days *in vitro* (DIV) shows that neurons treated with WT ACM have increased dendrite outgrowth relative to minimal media. Treating neurons with WT ACM with added SEMA3C is inhibitory to dendrite outgrowth in cortical neurons (Figure 1D).

**Figure 1.**
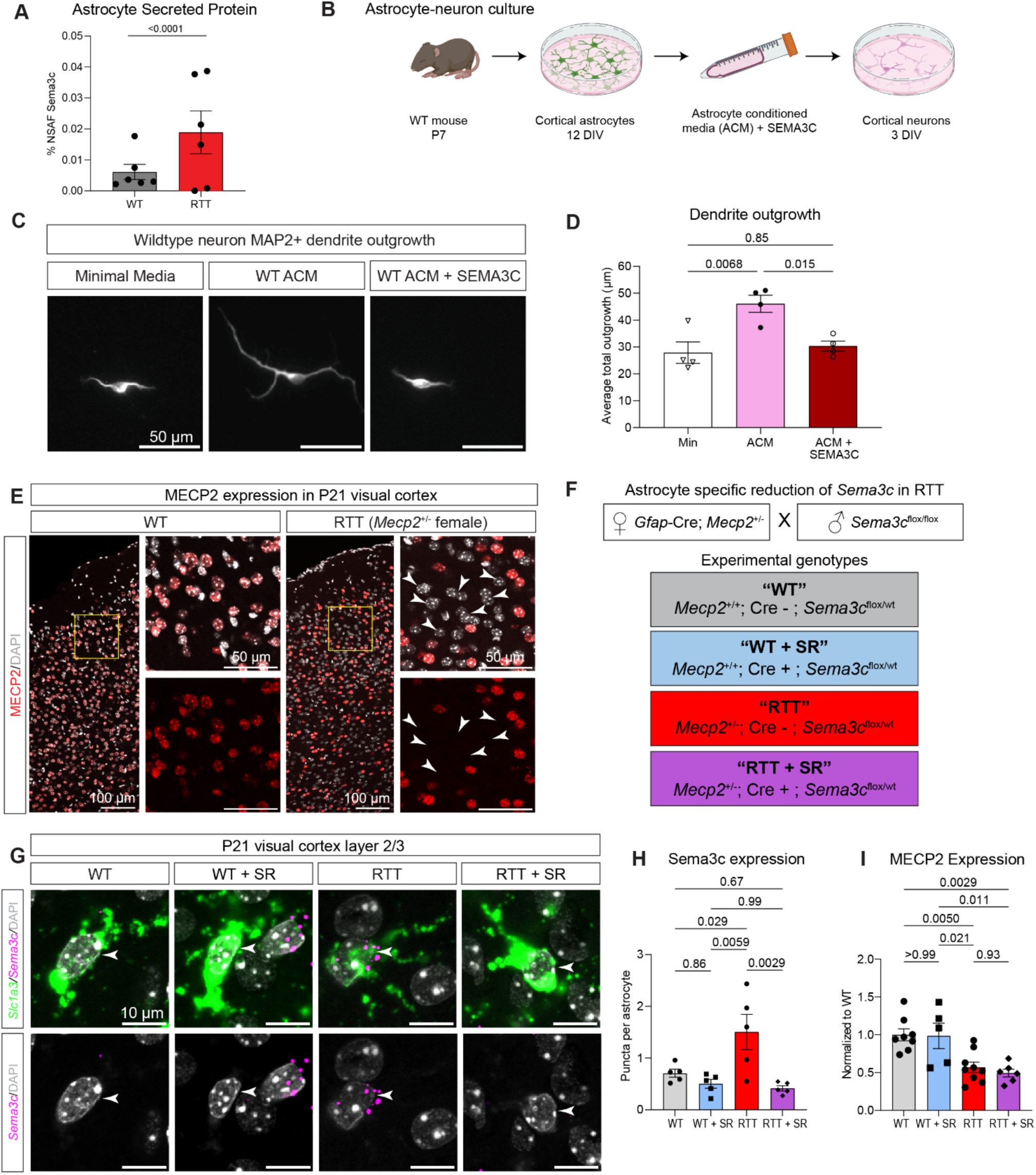
SEMA3C inhibits dendrite outgrowth and *in vivo* reduction strategy. **(A)** Quantification of % NSAF SEMA3C in astrocyte conditioned media from cultured wildtype (gray) and RTT (red) mouse model astrocytes (Mass Spectrometry: 0.0061 ± 0.0024 NSAF in WT ACM, 0.019 ± 0.070 NSAF in RTT ACM). Data from Caldwell et al., 2022. N = 6 cultures/group. Statistics by Tfold test in PatternLab (P <0.05, abundance > 0.01%, fold change between wildtype and RTT ≥ 1.5). **(B)** Workflow for measuring dendrite outgrowth. **(C)** Representative images of wildtype cortical dendrite outgrowth labeled by MAP2 with minimal media (left), WT ACM (middle), and WT ACM with added recombinant SEMA3C (right, 500ng/ml). Scale bars are 50 µm. **(D)** Quantification of average total dendrite outgrowth of neurons grown with minimal media (min, white), WT ACM (ACM, pink), or WT ACM + SEMA3C (dark red). N = 4 independent experiments with an average of 61 neurons per condition per replicate. One-way ANOVA with Tukey’s multiple comparisons. P values denoted. Error bars show SEM**. (E)** Representative images of MECP2 expression at P21 in WT or RTT (*Mecp2* heterozygous) females. For each genotype, left image shows a tile scan of the visual cortex (VC) (scale bar = 100 µm) with DAPI (gray) and MECP2 (red), the right zoomed in images (scale bar = 50 µm, location in tile scan denoted by yellow box) show DAPI and MECP2 (top) and MECP2 alone (bottom). Arrow heads label example DAPI nuclei that are MECP2 negative in RTT. **(F)** Breeding strategy for generating mice with astrocyte-specific reduction of SEMA3C (top). Experimental genotypes (bottom) are WT (gray), WT + SR (blue), RTT (red), and RTT + SR (purple). SR = SEMA3C reduction. **(G)** Representative images of P21 smFISH in VC L2/3 with (top) *Slc1a3* (green, astrocyte marker, alias GLAST), *Sema3c* (magenta) and DAPI (gray) and (bottom) *Sema3c* and DAPI, for each genotype. Scale bar = 10 µm. Arrowhead denotes *Slc1a3*-positive astrocyte nucleus. **(H)** Quantification of astrocyte *Sema3c* expression across genotypes shown as average number of *Sema3c* puncta per astrocyte. N = 5 mice per genotype. Each dot represents one animal. **(I)** Quantification of MECP2 expression across genotypes, normalized to WT from the same immunostaining experiment. N = 8 WT, 5 WT + SR, 9 RTT, and 6 RTT + SR animals. Each dot represents one animal. H-I. Two-way ANOVA with Tukey’s multiple comparisons. P values denoted. Error bars show SEM.

Since reduced dendrite outgrowth and arborization are phenotypic hallmarks of RTT, we tested the hypothesis that increased astrocyte secretion of SEMA3C contributes to RTT phenotypes. Our studies use *Mecp2* heterozygous^92^ females, which show mosaic expression of MECP2 (Figure 1E), and focus on the visual cortex, which shows changes in neuronal morphology and activity in RTT^31,46,47,93–95^. To normalize SEMA3C levels specifically in astrocytes, *Mecp2* heterozygous females were first crossed with GFAP-Cre73.12^100,10^ mice and the offspring were then crossed with *Sema3c* floxed mice^96^ (Figure 1F). This conditionally deletes one *Sema3c* allele in astrocytes and reduces gene dosage to wildtype physiological levels rather than eliminating expression entirely. In each experiment, we examine four genotypes: wildtype (WT), wildtype with SEMA3C reduction (WT + SR), RTT, and RTT with SEMA3C reduction (RTT + SR). To validate reduction of SEMA3C, we used single molecule fluorescent *in situ* hybridization (smFISH) at P21 for *Sema3c* mRNA transcripts in *Slc1a3+* astrocytes (Figure 1G). We find that *Sema3c* mRNA transcript abundance is increased in astrocytes in RTT relative to WT. RTT + SR reduces *Sema3c* in astrocytes, relative to RTT, and normalizes to WT levels (Figure 1H). Astrocyte SEMA3C reduction does not impact MECP2 expression (Figure 1I). Thus, *Sema3c* is increased in RTT, and RTT + SR mice provide a model to investigate the functional consequences of this.

### Astrocyte SEMA3C reduction improves dendritic arborization in RTT

Given that reduced dendritic arborization is reported in human postmortem RTT tissue^41,42^ and in cortical neurons of RTT mouse models^49,77,93^, and that SEMA3C is sufficient to inhibit dendrite outgrowth in cell culture (Figure 1D), we hypothesized that reduced dendritic arborization in RTT is mediated by increased astrocyte SEMA3C. To examine this, we labeled individual visual cortex L2/3 pyramidal neurons at P60 (Figure 2A) and quantified their dendrite arborization using Sholl analysis (Figure 2B). We found that RTT neurons have reduced dendrite arborization relative to WT. Notably, RTT + SR normalizes arborization suggesting that dendritic outgrowth is inhibited by astrocyte-derived SEMA3C *in vivo*. To compare dendrite arborization in RTT neurons that are MECP2 positive versus negative, we conducted antibody staining for MECP2 protein (Figures S2A and S2B). MECP2 positive and MECP2 negative RTT neurons do not show significantly different dendritic arborization (Figure S2C), suggesting that MECP2 expression does not exert a cell autonomous effect on dendritic arborization in L2/3 visual cortex pyramidal neurons.

**Figure 2.**
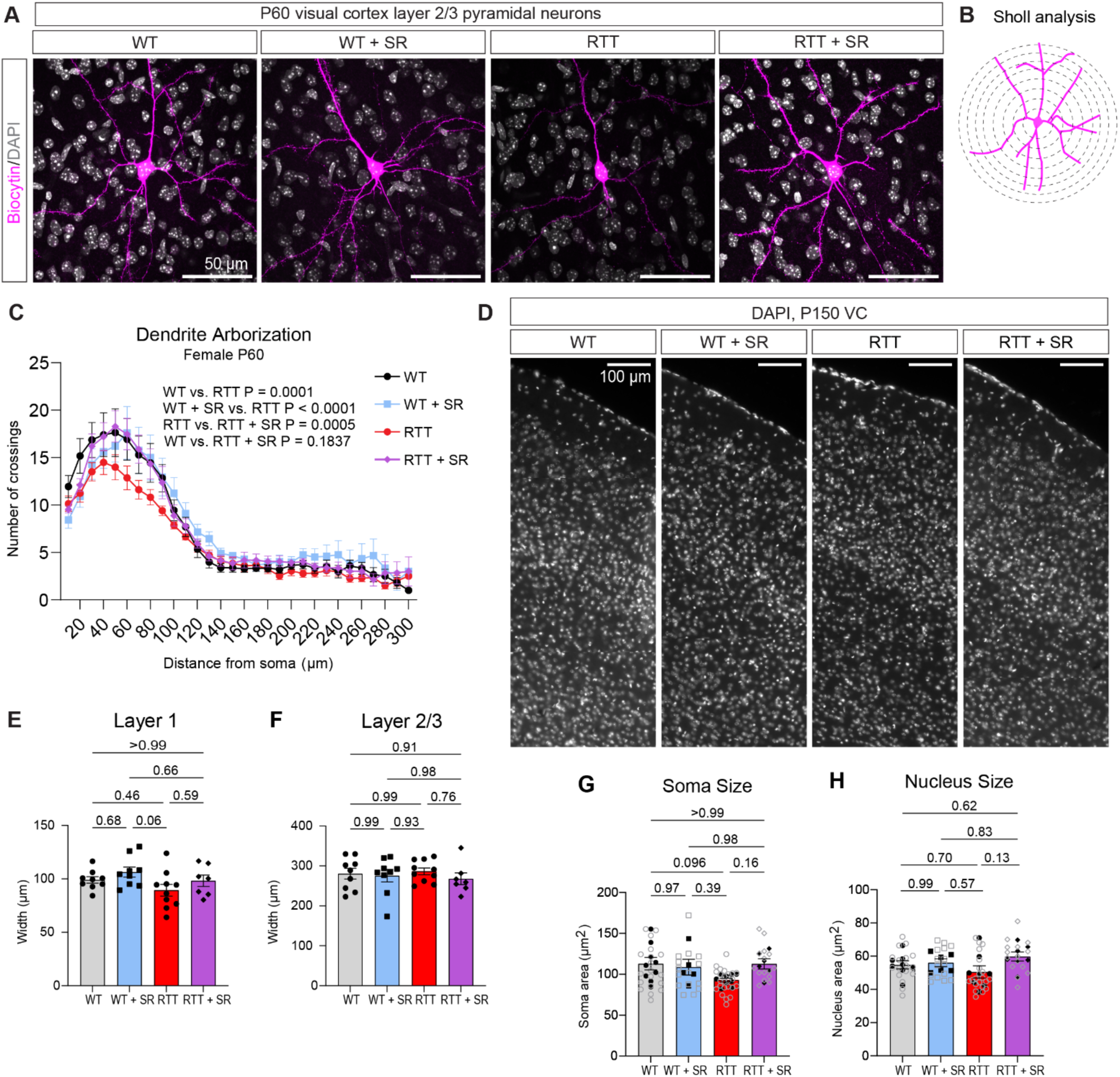
Astrocyte SEMA3C reduction improves dendrite arborization in Rett syndrome. **(A)** Representative images of individual VC L2/3 pyramidal neurons (biocytin filled neuron in magenta, DAPI in gray). Scale bar is 50 µm. (**B)** Dendrite arborization is analyzed using Sholl analysis to measure dendrite branching points at 10 µm increments from the soma. (**C)** Quantification of dendrite arborization. Mixed-effects model (REML), two factors: genotype and distance from soma; Tukey’s multiple comparisons. P values denoted on figure. (**D)** Representative images of VC with DAPI (gray). Scale bar is 100 µm. (**E)** Quantification of VC layer 1 width at P150. **(F)** Quantification of VC L2/3 width at P150. E-F. N = 10 WT, 9 WT + SR, 10 RTT, and 7 RTT + SR. Each symbol represents one animal. **(G)** Quantification of soma area. **(H)** Quantification of nucleus area. Closed black symbols represent animals while open gray symbols represent individual cells. G-H. N = 8 WT, 5 WT + SR, 9 RTT, and 6 RTT + SR animals with 1-4 neurons per animal. E-H. Two-way ANOVA on per-mouse means with Tukey’s multiple comparisons. P values denoted. Error bars depict SEM.

Region-specific cortical volume and cortical layer width reductions occur in individuals with RTT and in mouse models^48,97–103^. Therefore, we measured the width of layers 1 and 2/3 of the visual cortex. We observe no difference in width of either layer in WT versus RTT females (Figures 2D-2F). We also found no significant difference in L2/3 pyramidal neuron soma or nucleus size (Figures 2G and 2H), although WT vs. RTT comparison for soma size approaches significance (P = 0.096). Taken together, we find limited RTT effects on cortical layer width or neuronal nucleus size in the female *Mecp2* heterozygous visual cortex, but dendrite arborization is significantly diminished. Our data reveals that reducing astrocyte SEMA3C can improve dendritic arborization deficits in RTT, suggesting that increased astrocyte SEMA3C secretion in RTT contributes to neuronal morphology deficits.

### Reduction of astrocyte SEMA3C increases spine density

Postmortem tissue from individuals with RTT and mouse models of RTT show reduced dendritic spine density and altered spine morphology^42,45,47,77,104^. To investigate if spine density and morphology are altered in the female RTT visual cortex, and to ask if SEMA3C reduction can impact this, we examined spines of P60 L2/3 pyramidal neurons. Using high resolution confocal imaging, we imaged basal and apical dendritic spines of WT, WT + SR, RTT, and RTT + SR neurons. We analyzed primary basal dendrites and the distal segments of apical dendrites, each at approximately the same distance from the soma. Both basal and apical spines (Figures 3A-3D; S2A and S2B) on RTT neurons do not show altered spine density relative to WT. We did, however, observe that SEMA3C reduction increases spine density in both WT and RTT genotypes (Basal spines: two-way ANOVA SEMA3C effect P = 0.0091, Figure 3B. Apical spines: two-way ANOVA SEMA3C effect P = 0.0022, Figure 3D).

**Figure 3.**
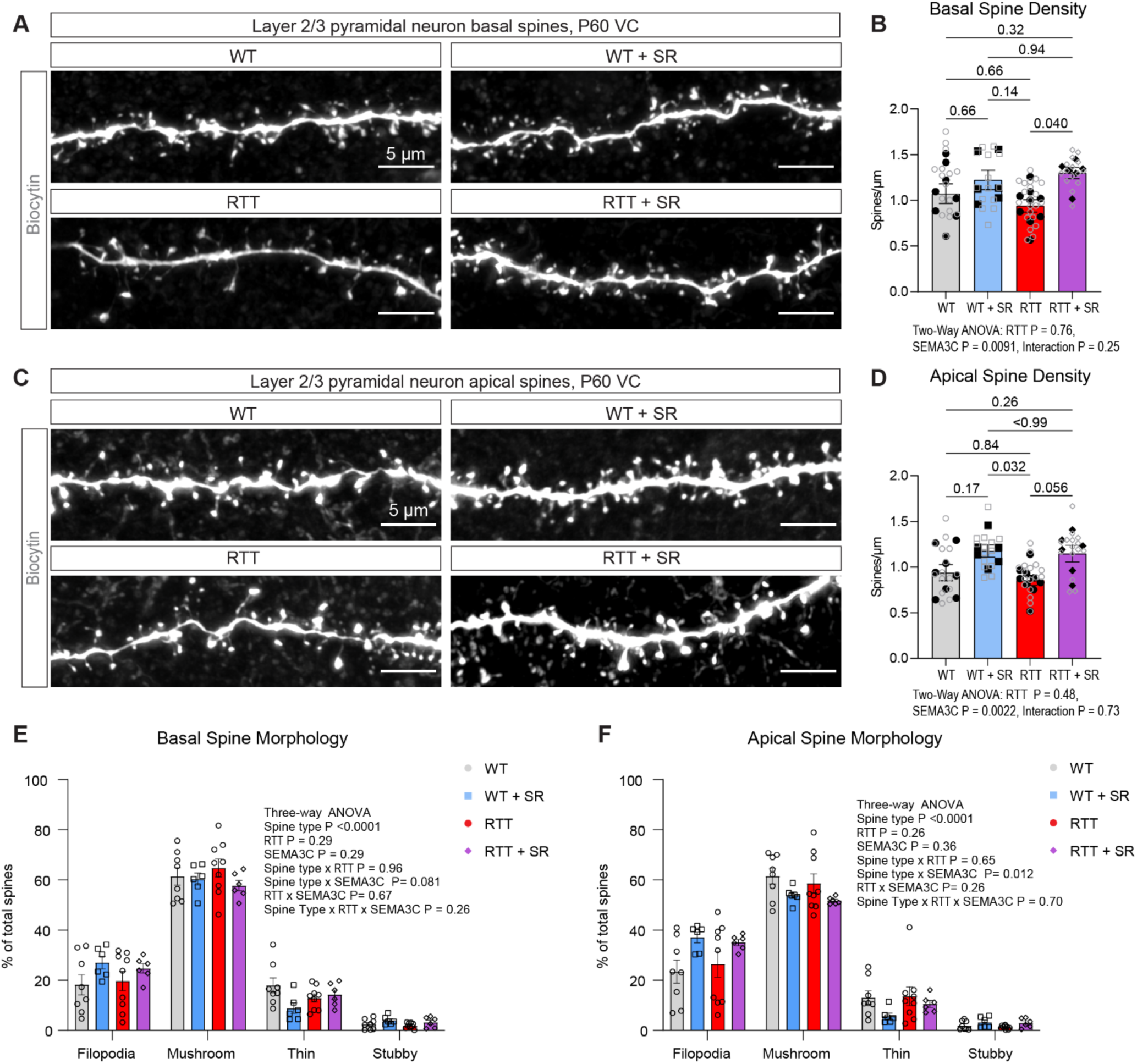
Reduction of astrocyte SEMA3C increases spine density. **(A)** Representative images of L2/3 pyramidal neuron basal dendritic spines. Scale bar = 5 µm. **(B)** Quantification of basal spine density (number of spines per µm). Two-way ANOVA with Tukey’s multiple comparisons. P values denoted. **(C)** Representative images of L2/3 pyramidal neuron apical dendritic spines. Scale bar = 5 µm. **(D)** Quantification of apical spine density. Two-way ANOVA with Tukey’s multiple comparisons. P values denoted. B, D. Closed black symbols represent animals while open gray symbols represent individual cells. Statistics are conducted on animal averages. **(E-F)** Quantification of spine morphology showing the percentage of spines that are filopodia, mushroom, thin, or stubby morphology for (E) basal and (F) apical spines. Each symbol represents one animal. Three-way ANOVA on per-mouse means. P values denoted. B, D, E, F. N = 8 WT, 6 WT + SR, 9 RTT, and 6 RTT + SR animals. For basal dendrites: 16 WT neurons, 14 WT + SR neurons, 21 RTT neurons, and 15 RTT + SR neurons. For apical dendrites: 15 WT neurons, 14 WT + SR neurons, 17 RTT neurons, and 15 RTT + SR neurons. 5 apical and basal dendrites were analyzed for each neuron. Error bars show SEM.

We next classified spines based on morphology (filopodia, mushroom, thin, or stubby)^105^, finding spine morphology is not altered in RTT neurons overall, but that there is a spine morphology x SEMA3C interaction effect in apical dendrites of both WT and RTT neurons (Figures 3E-3F). Comparison of MECP2-positive and MECP2-negative neurons in RTT mice also revealed no difference in spine density, indicating that MECP2 expression does not have a cell-autonomous effect on this parameter (Figures S2C-S2D). In contrast, MECP2 expression cell-autonomously impacts spine morphology, as neurons lacking MECP2 in RTT mice showed an increased proportion of filopodia-type spines and a decreased proportion of mushroom-type spines compared to neurons expressing MECP2 (Figures S2E-S2F). Overall, we find that neurons in RTT mice show minimal alterations in spine density and spine morphology in visual cortex L2/3 pyramidal neurons at P60, and that reducing astrocytic SEMA3C is sufficient to increase spine density independently of a RTT phenotype.

### Astrocyte SEMA3C reduction in RTT normalizes sIPSC frequency

We next examined synaptic activity in female RTT model mice and tested whether astrocyte-specific SEMA3C reduction altered these electrophysiological phenotypes. Previous cellular electrophysiological analyses of RTT model mice have primarily focused on males while our study focuses on females^43,46,79,106^. In male *Mecp2-*null visual cortex, neuronal activity propagation to upper cortical layers is abnormally suppressed at P22-25^95^, while at P45, visually evoked excitatory and inhibitory synaptic events onto L2/3 pyramidal neurons are both reduced^46^. Together, these studies establish visual cortex circuit dysfunction in male RTT models, but how synaptic activity is altered in female RTT model mice remains less well understood. Here, we conducted *ex vivo* whole-cell patch clamp recordings from L2/3 pyramidal neurons at P60 (Figure 4A), corresponding to the age of disease onset^74,78^. We recorded spontaneous excitatory and inhibitory postsynaptic currents (sEPSCs and sIPSCs, respectively) (Figures 4B-4C). Analysis of sEPSCs revealed no significant differences in average frequency across groups (Figures 4D and S3A). Average sEPSC amplitude does not differ between WT and RTT mice, but there was a significant effect of SEMA3C reduction, towards lower sEPSC amplitudes in SEMA3C-reduced groups (Figures 4E and S3B).

**Figure 4.**
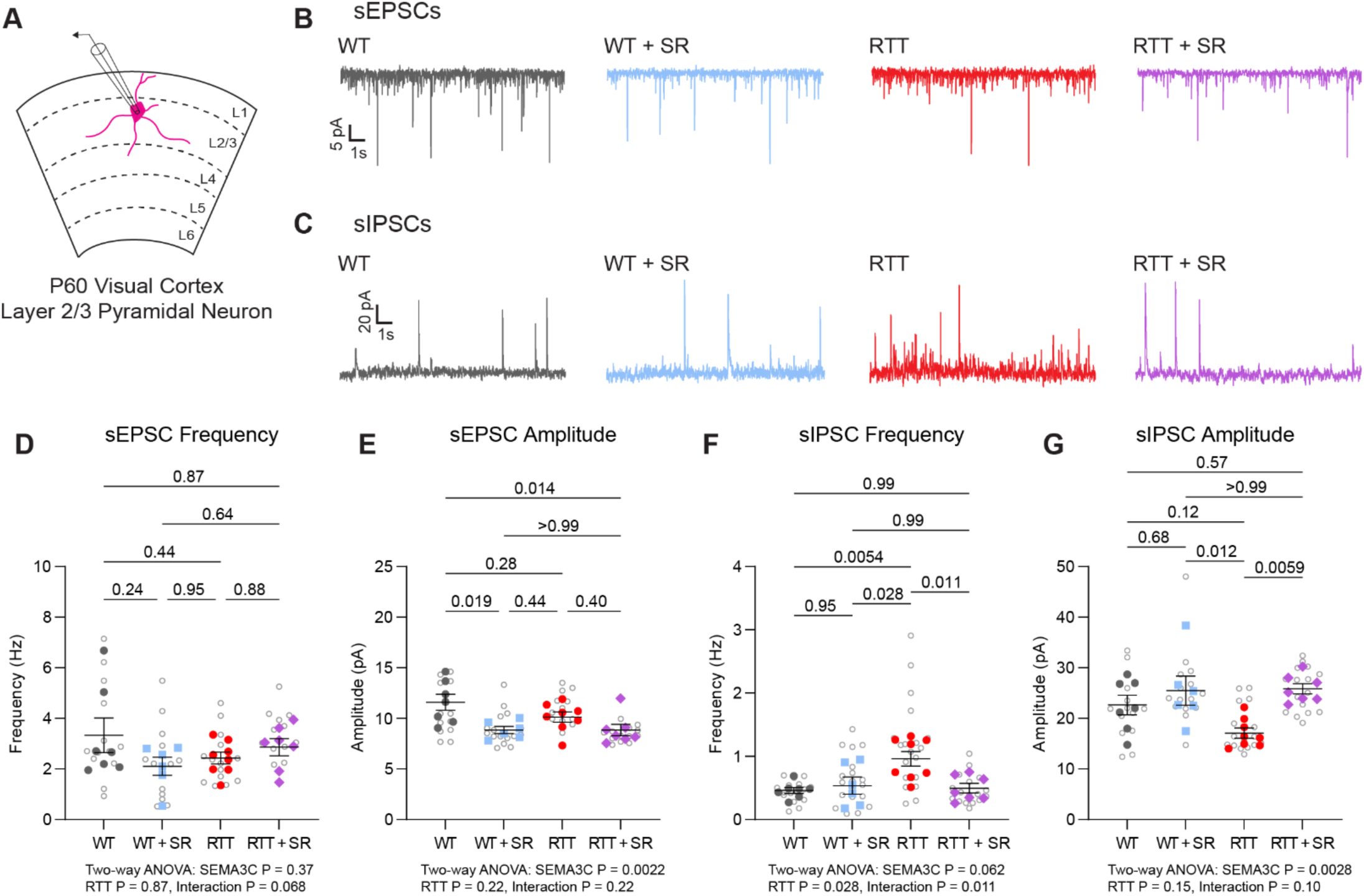
Astrocyte SEMA3C reduction in Rett syndrome normalizes sIPSC frequency. **(A)** Schematic depicting whole-cell patch clamp recording from VC L2/3 pyramidal neurons at P60. **(B)** Representative excitatory spontaneous postsynaptic currents (sEPSCs) from each genotype. **(C)** Representative inhibitory spontaneous postsynaptic currents (sIPSCs) from each genotype. **(D)** sEPSC frequency (Hz). **(E)** sEPSC amplitude (pA). **(F)** sIPSC frequency (Hz). **(G)** sIPSC amplitude (pA). D-G. Gray open circles represent individual cells. Black, blue, red, or purple solid points represent animal averages. N = 7 WT, 6 WT + SR, 8 RTT, and 7 RTT + SR animals with 1-4 cells per animal. Two-way ANOVA on per-mouse means with Tukey’s multiple comparisons. P values denoted. Error bars depict mean ± SEM.

Notably, sIPSC frequency is significantly increased in RTT mice relative to WT. This phenotype is normalized to WT levels in RTT + SR mice (Figures 4F and S3C). Thus, spontaneous inhibitory event frequency is elevated in female RTT mice and SEMA3C reduction can normalize this activity. sIPSC amplitude is not significantly different between WT and RTT neurons, although RTT neurons show lower mean amplitudes. SEMA3C reduction increases sIPSC amplitude overall, with RTT + SR amplitudes significantly higher than RTT and not significantly different from WT (Figures 4G and S3D).There are no differences in resting membrane potential (Figure S3E). Together, these results identify increased spontaneous inhibitory event frequency as a prominent electrophysiological phenotype in female RTT visual cortex that is normalized by astrocyte SEMA3C reduction.

### Astrocyte SEMA3C reduction rescues RTT behavioral deficits

Given improvements in neuronal structure and synaptic activity, we next asked if astrocyte *Sema3c* reduction can improve RTT behavioral deficits, focusing on visual acuity, motor, and anxiety-like behavior (Figure 5A), which are altered in female RTT mouse models^52,78,80,92,95,107^ and individuals with RTT^31,108,109^. We observed that mouse litters have the expected inheritance pattern with no skew or loss of one genotype (Figure S4A). We found that RTT body weight at P120 is increased relative to WT (Figure S4B) as others have reported^78,85^. RTT + SR body weight does not differ significantly from WT or RTT suggesting an intermediate phenotype. To measure visual acuity, we used the non-invasive optomotor assay which measures the involuntary responses of mice to visual stimuli^110^. In line with previous reports^31,95^, visual acuity is reduced in RTT relative to WT mice. Importantly, we found that RTT + SR rescues visual acuity to WT levels (Figure 5B) indicating that increased astrocyte SEMA3C contributes to visual deficits in RTT. Astrocyte SEMA3C reduction improves visual acuity even in male *Mecp2*-null mice at P45 (Figure S4C), highlighting that RTT + SR can still rescue visual deficits despite the more severe phenotype.

**Figure 5.**
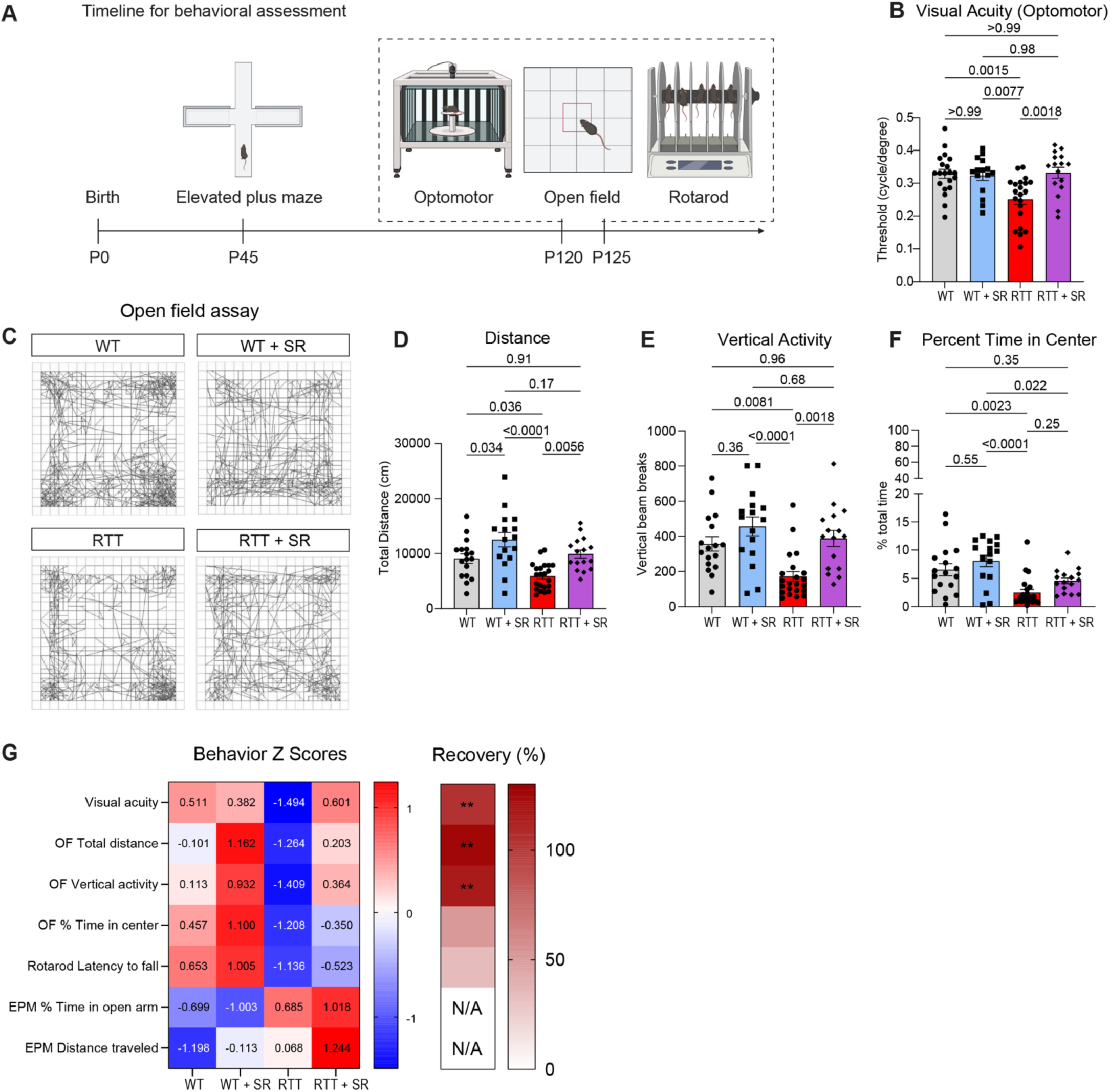
Astrocyte SEMA3C reduction in Rett syndrome rescues visual acuity and open field activity. **(A)** Timeline for behavioral assessment. **(B)** Quantification of visual acuity threshold (cycles/degree) in the optomotor assay. N = 20 WT, 15 WT + SR, 21 RTT, and 16 RTT + SR mice. **(C)** Representative traces for open field horizontal activity. **(D-F)** Quantification of open field activity over the 60-minute assay: (D) total distance (cm), (E) vertical activity (rearing), and (F) percentage of time spent in center of the open field arena. N = 17 WT, 16 WT + SR, 22 RTT, and 16 RTT + SR mice. **(G)** Heatmap of behavioral Z-scores across genotypes. OF = open field. EPM = elevated plus maze. The percent recovery heatmap shows the average percent recovery of RTT + SR towards WT levels (RTT + SR – RTT)/(WT – RTT) x 100. B, D-F. Two-way ANOVA with Tukey’s multiple comparisons. P values denoted. Error bars show SEM. Each dot represents one mouse.

To test motor behavior, we employed the rotarod and open field assays. In the rotarod assay, a measure of motor coordination and balance, RTT mice show reduced latency to fall, which is not improved in RTT + SR mice (Figure S4D and S4E). In the open field assay, both distance traveled and vertical activity (rearing) are reduced in RTT mice relative to WT, while RTT + SR normalizes these behaviors to WT levels (Figures 5C-5E). Together, these findings suggest that astrocyte SEMA3C reduction improves spontaneous locomotor and exploratory activity, but not rotarod performance, which requires additional coordination and balance demands. WT mice with SEMA3C reduction exhibited increased activity in the open field assay, suggesting that astrocyte SEMA3C may contribute to typical motor behavior.

To examine anxiety-related behavior we also analyzed the percentage of time spent in the center of the arena in the open field assay^111^. We found a significant reduction in the percentage of time spent in the center for RTT mice which may suggest increased anxiety-like behavior (Figure 5F). The percentage of time spent in the center by RTT + SR mice does not significantly differ from WT or RTT mice suggesting an intermediate phenotype. We also examined anxiety-like behavior in the elevated plus maze (EPM), at P45 prior to the onset of motor deficits (Figures S4F-S4H). We did not observe robust WT versus RTT differences in the EPM. Overall, these results indicate that RTT + SR can restore two different key behavioral deficits in RTT: visual acuity and open field activity (Figure 5G). Taken together with our neuron morphology and synaptic activity analyses, we find astrocyte SEMA3C contributes to RTT deficits across cellular and functional levels.

### SEMA3C signals through PLXND1 and NRP2 to inhibit dendrite outgrowth in wildtype neurons

To investigate the mechanism of SEMA3C signaling to neurons, we next asked which receptor SEMA3C signals through in P7 wildtype cortical neurons. Semaphorins interact with neuropilin and plexin receptors. Neuropilins are transmembrane glycoproteins with short cytoplasmic domains that lack intracellular signaling capabilities^67^ while plexins are co-receptors that mediate intracellular signaling via an intracellular GTPase activating protein domain^68^. Within these protein classes, Plexin D1 (PLXND1), Neuropilin-1 (NRP1), and (Neuropilin-2) NRP2 are known SEMA3C receptors^90,112–114^ (Figure 6A) and are expressed in mouse cortical neurons^115^ (Figure S5A). To test if these receptors mediate the inhibitory effects of SEMA3C, we used cortical neurons cultured with astrocyte conditioned media (ACM) to promote dendrite outgrowth, along with SEMA3C to inhibit this, as described in Figure 1. We applied receptor-specific blocking antibodies against PLXND1, NRP1, or NRP2 in the presence of SEMA3C and ACM and then quantified dendrite outgrowth after three days *in vitro*. We found that blocking PLXND1 increases dendrite outgrowth relative to SEMA3C alone or with an IgG control (Figures 6C-6D and S5B). Likewise, blocking NRP2 increases dendrite outgrowth, relative to SEMA3C or the IgG control (Figures 6E-6F and S6C). Since blocking PLXND1 or NRP2 resulted in similar outgrowth to adding WT ACM alone, this suggests that blocking either receptor is sufficient to prevent SEMA3C inhibition of dendrite outgrowth in WT neurons. In contrast, blocking NRP1 does not improve outgrowth (Figures 6G-6H and S5D), indicating SEMA3C does not mediate its negative effect on dendritic outgrowth through NRP1. In summary, blocking PLXND1 or NRP2 reduces SEMA3C’s inhibitory effects on dendrite outgrowth in wildtype neurons.

**Figure 6.**
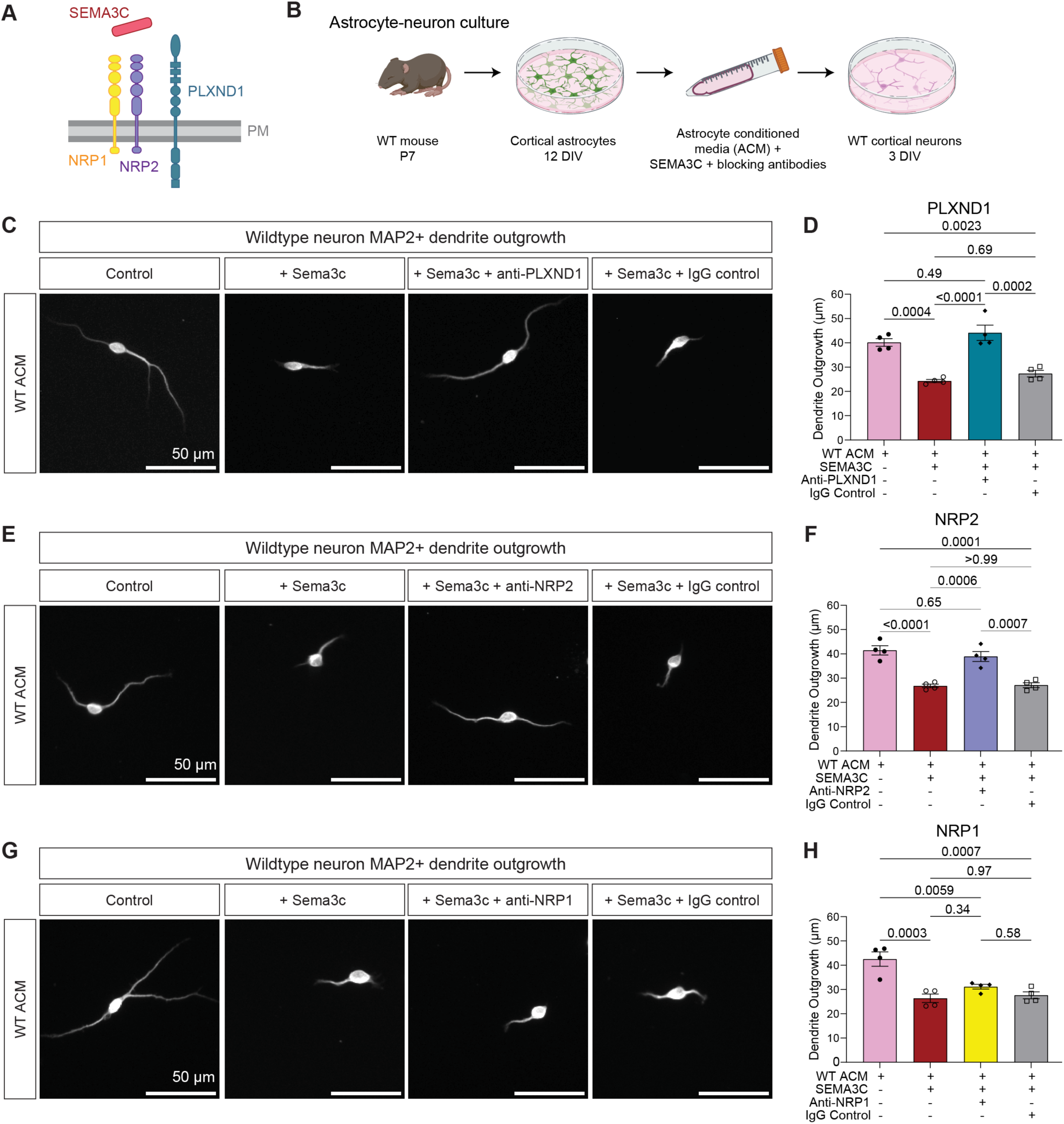
SEMA3C inhibits dendrite outgrowth through PLXND1 and NRP2 receptors in wildtype neurons. **(A)** Schematic depicting candidate SEMA3C receptors in the plasma membrane (PM). **(B)** Workflow for testing blocking antibody impact on SEMA3C inhibition of dendrite outgrowth. **(C, E, G)** Representative images of WT neuron dendrite outgrowth with WT ACM alone (control), with added recombinant SEMA3C (200ng/ml), with SEMA3C and blocking antibody (C. 5µg/ml anti-PLXND1, E., 10µg/ml anti-NRP2 and G. 10µg/ml anti-NRP1), or with SEMA3C and IgG control (C. 5µg/ml, E, G. 10µg/ml). **(D, F, H)** Quantification of average total dendrite outgrowth for ACM alone, ACM with SEMA3C, ACM with SEMA3C with blocking antibody, and ACM with SEMA3C and IgG control. N = 4 independent experiments with an average of 77 neurons per condition per replicate. C, E, G. Scale bars are 50 µm. D, F, H. One-way ANOVA with Tukey’s multiple comparisons. P values denoted. Error bars show SEM.

### Blocking PLXND1 improves dendrite outgrowth in the presence of RTT ACM

ACM from RTT astrocytes has been shown to inhibit dendrite outgrowth^15,17^, so we next asked if upregulated SEMA3C in RTT ACM is contributing to this inhibition by signaling through the identified receptors. To address this, we asked whether blocking PLXND1 or NRP2 receptors in cortical neurons treated with RTT ACM would be sufficient to improve dendrite outgrowth, which we tested by applying RTT ACM to neurons alongside receptor-specific blocking antibodies. To account for the presence of both MECP2-positive and MECP2-negative neurons in the female *Mecp2* heterozygous brain, we tested RTT ACM effects on both wildtype neurons and MECP2-negative RTT neurons derived from *Mecp2*-null male mice (Figures 7A-7C). We found that blocking PLXND1 (Figure 7D and S6A) or NRP2 (Figures 7E and S6B) in RTT ACM improved dendrite outgrowth relative to RTT ACM alone or with IgG control in wildtype neurons. Blocking NRP1 did not improve wildtype dendrite outgrowth in the presence of RTT ACM (Figures 7F and S6C), as also observed in the presence of SEMA3C. In RTT neurons, blocking PLXND1 (Figures 7G and S6D), but not NRP2 or NRP1 (Figures 7H-7I and S6E-S6F) improved dendrite outgrowth, demonstrating a genotype-dependent effect. SEMA3C can also signal through PLXNA2^116,117^, which is expressed by visual cortex layer 2/3 glutamatergic neurons^115^, however blocking PLXNA2 in the presence of RTT ACM did not improve dendrite outgrowth in either wildtype or RTT neurons (Figures S7G-S7L). Thus, in wildtype neurons, blocking PLXND1 or NRP2 improves dendrite outgrowth in the presence of recombinant SEMA3C or RTT ACM, whereas in RTT neurons only blocking PLXND1 improves outgrowth. Altogether, our findings reveal elevated astrocyte SEMA3C as a mechanism through which astrocytes contribute to altered neuronal morphology, synaptic function, and behavior in RTT.

**Figure 7.**
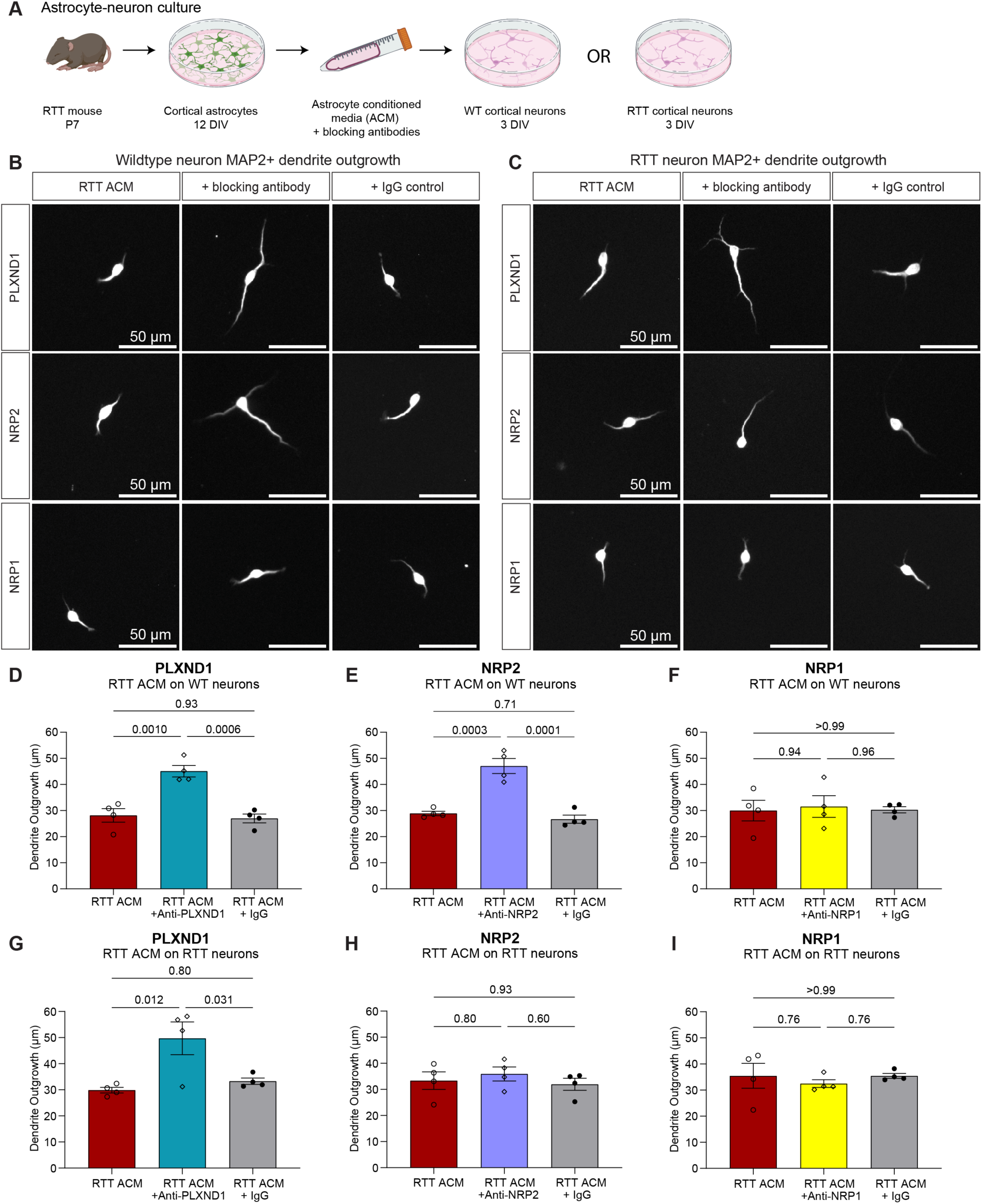
SEMA3C inhibits dendrite outgrowth through PLXND1 receptor in RTT neurons. **(A)** Workflow for testing blocking antibody impact on RTT astrocyte conditioned media inhibition of dendrite outgrowth in WT or RTT neurons. RTT astrocytes and neurons were derived from *Mecp2*-null male mice and therefore lack MECP2. **(B-C)** Representative images of (B) WT neuron dendrite outgrowth or (C) RTT neuron dendrite outgrowth with RTT ACM (left panels), RTT ACM with blocking antibodies (middle panels) to PLXND1 (top row, 5µg/ml), NRP2 (middle row, 10µg/ml)m or NRP1 (bottom row, 10µg/ml), and IgG control (right panels, 5µg/ml for PLXND1 control, 10µg/ml for NRP1/2 control). Scale bars are 50 µm. **(D-F)** Quantification of average total dendrite outgrowth for RTT ACM alone, with blocking antibody (D. 5µg/ml anti-PLXND1, E., 10µg/ml anti-NRP2 and F. 10µg/ml anti-NRP1), or with IgG control (D. 5µg/ml, E., F. 10µg/ml) on WT neurons. **(G-I)** Quantification of average total dendrite outgrowth for RTT ACM alone, with blocking antibody (G. 5µg/ml anti-PLXND1, H., 10µg/ml anti-NRP2 and I. 10µg/ml anti-NRP1), or with IgG control (G. 5µg/ml, H., I. 10µg/ml) on RTT neurons. N = 4 independent experiments with an average of 64 neurons per condition per replicate. C, E, G. Scale bars are 50 µm. D, F, H. One-way ANOVA with Tukey’s multiple comparisons. P values denoted. Error bars show SEM.

## Discussion

In this study we show that astrocyte SEMA3C is elevated *in vivo* in RTT, and that normalization of astrocyte SEMA3C to WT physiological levels improves deleterious phenotypes in female RTT mice at multiple levels of analysis, from cellular to behavioral. By studying female *Mecp2* heterozygous mice, we tested the impact of astrocyte SEMA3C reduction in a *Mecp2* mosaic model that reflects an important feature of RTT biology. We demonstrate that RTT neuronal morphology and synaptic activity deficits can be improved through astrocyte specific reduction of SEMA3C. Visual acuity and open field activity were also restored to WT levels, demonstrating functional improvement. Consistent with the reduced dendritic arborization observed in RTT, we find that SEMA3C is inhibitory to dendrite outgrowth through neuronal PLXND1 receptor signaling. Thus, dysregulated SEMA3C-PLXND1 signaling contributes to RTT neuroanatomical and behavioral deficits. This work highlights astrocyte-secreted factors as active contributors to neuronal development and function whose dysregulation can shape neurodevelopmental disorder phenotypes.

A key finding of this study is that astrocyte SEMA3C reduction improves dendritic complexity in RTT. Altered dendritic arborization is a shared feature across multiple neurodevelopmental disorders, including Down syndrome, Fragile X syndrome, and other syndromic forms of autism spectrum disorders and intellectual disability^118,119–123^. Since astrocyte SEMA3C secretion is increased across RTT, Fragile X syndrome, and Down syndrome^12^, dysregulated astrocyte semaphorin signaling may represent a convergent pathway contributing to altered dendritic architecture across distinct neurodevelopmental disorders. This raises the broader possibility that altered astrocyte-secreted signals contribute to shared features of neuronal dysfunction.

Previous studies in *Mecp2-*null male models reported reductions in cortical layer width and soma size^48,102^, while studies in the motor cortex of *Mecp2* heterozygous females identified cell-autonomous effects of MECP2 expression on neuron morphology^49^. In our study, we did not detect significant reductions in cortical layer or soma size in the female RTT visual cortex, nor did we find significant cell autonomous effects of MECP2 expression on soma size or dendritic arborization. These differences may reflect region-specific phenotypes, the female mosaic context, or non-cell autonomous influences from neighboring MECP2-positive cells. However, our sample size of MECP2-positive neurons was limited, and future studies should further investigate how MECP2 mosaicism impacts visual cortex phenotypes. Likewise, we did not detect significant reductions in spine density at P60, although later time points may reveal changes similar to those reported in the MECP2 heterozygous motor cortex^77^. Notably, MECP2-negative RTT neurons showed increased filopodia-shaped spines, consistent with prior reports of immature spine morphology in RTT^42,44,45^.

Our electrophysiological analysis of visual cortex neurons identifies elevated sIPSC frequency as a prominent synaptic phenotype in female RTT mice that is normalized by astrocyte SEMA3C reduction. This finding indicates that astrocyte-secreted SEMA3C contributes to abnormal inhibitory synaptic activity in RTT. The mechanisms linking SEMA3C signaling to changes in synaptic activity remain to be determined. Semaphorin pathways have been implicated in synaptic regulation, including SEMA3F-NRP2-PLXNA3^119^ and SEMA4A/D-PLXNB1^120–122^ regulation of excitatory and inhibitory postsynaptic receptors, respectively. One possibility is that astrocyte SEMA3C interaction with neuronal PLXND1 receptors alters synaptic receptor organization or function in a manner that changes inhibitory synaptic activity. Additionally, increased SEMA3C signaling may impact inhibitory neuron function directly.

To understand if RTT + SR could improve RTT behavioral deficits, we conducted behavioral assessments known to be altered in female RTT mouse models and with construct validity for human motor and visual acuity deficits in RTT^31,108,109^. We find improvement of visual acuity and motor behavior in the open field assay when astrocyte SEMA3C is reduced in RTT mice. We did not find behavioral improvement in the rotarod assay which could reflect deficits in motor coordination, muscle strength, or both^78,123^. Thus, the beneficial effects of reducing astrocyte SEMA3C are behavior-specific rather than global. Our proteomic^12^, cell culture assays, and histological analysis focused on the cortex while motor coordination also engages noncortical circuitry. Behavioral phenotypes not restored by astrocyte SEMA3C reduction may reflect the contributions of additional astrocyte-derived signals altered in RTT.

We find that both SEMA3C and RTT astrocyte conditioned media inhibit dendrite outgrowth through PLXND1-dependent signaling. In wildtype neurons, NRP2 blockade can also improve outgrowth. PLXND1 is an R-Ras GTPase activating protein. Semaphorin signaling activates the plexin GAP domain to downregulate R-Ras activity and inhibits neurite outgrowth^124–130^, which may be one mechanism by which SEMA3C regulates dendrite outgrowth. PLXND1 and NRP2 knockout mice show changes to spine density, synaptic activity, and behavior^62,64,131,132^. NRP2 knockout mice have increased spine density suggesting NRP2 regulation of spines in line with the modest increase in spine density observed in mice with astrocyte SEMA3C reduction. Another semaphorin family member, SEMA3E, also shows increased astrocyte secretion in RTT, Fragile X syndrome, and Down syndrome^12^. Additional studies are needed to determine whether SEMA3E, a known PLXND1 ligand^133,134^, contributes to dendritic arborization, synaptic activity, and behavioral phenotypes. Semaphorin signaling family members are also implicated in autism spectrum disorders^135–137^, suggesting that astrocyte semaphorin signaling may be relevant across multiple neurodevelopmental disorders.

Taken together, this study finds that astrocyte secreted SEMA3C is a non-cell autonomous contributor to neuronal alterations in RTT syndrome. Reducing SEMA3C level in astrocytes restores morphological, functional, and behavioral deficits, likely in part through PLXND1 signaling. These findings identify astrocyte-secreted SEMA3C as a disease relevant regulator of neuronal structure and function in RTT and highlight astrocyte-neuron semaphorin signaling as a candidate therapeutic pathway for RTT and potentially other neurodevelopmental disorders.

## Resource availability

### Lead Contact

Correspondence and request for materials should be addressed to Nicola J. Allen.

## Acknowledgements

This work was supported by the Chan Zuckberberg Initiative and NIH R21NS139073 to N.J.A. K.A.L. was supported by NIH K00NS108515, NIH K99NS140545, and Burroughs Wellcome Fund 1287660. A.M. was supported by Colors of the Brain/Kavli Institute for Brain and Mind Scholars Program. This work was supported by the In Vivo Scientific Services Facility of the Salk Institute and the Waitt Advanced Biophotonics Core Facility of the Salk Institute with funding from NIH-NCI CCSG P30 CA014195, NIH-NIA San Diego Nathan Shock Center P30 AG068635, The Henry L. Guenther Foundation and the Waitt Foundation. We thank N. Andrews and K. Hitchcock for technical assistance with behavioral experiments; A. Calmont for the *Sema3c* floxed mice, J. Hash, M. Iyer, C. Chambers, Y. Nichols, and the Salk Animal Resources Department for assistance with the mouse colony; L. Orefice, L. Mamounas, E. Mulhall, and members of the Allen lab and Salk MNL for feedback and discussions. Biorender was used to create schematics.

## Author contributions

Conceptualization: K.A.L. and N.J.A. Experimental work: K.A.L., A.P., A.K., and A.M. Data analysis: K.A.L., A.P., N.J.A. Figure preparation and writing: K.A.L Editing: K.A.L., and N.J.A. Funding acquisition: K.A.L. and N.J.A.

## Methods

### Experimental model and subject details

#### Mice

All animal work was approved by the Salk Institutional Animal Care and Use Committee (IACUC). Mice were housed in the Salk Institute vivarium with ad libitum food and water access and a 12:12 light and dark cycle. All experiments used littermates as controls.

##### *Mecp2* mice

*Mecp2* mice (B6.129P2(C)-Mecp2tm1.1Bird/J, RRID: IMSR_JAX:003890) were maintained by crossing female *Mecp2* heterozygous mice to C57/BL6 males.

##### *Sema3c* flox mice

*Sema3c* flox mice were obtained from Dr. Amelie Calmont originally generated through the Mouse Biology Program at UCD using a targeting vector from the Wellcome Trust Sanger Institute (project ID 69993, MGI:107557). *Sema3c* flox mice were maintained with a C57/BL6 background.

##### GFAP-Cre mice

GFAP-Cre 73.12 (Jax: 012886) were maintained by crossing Cre positive females to C57/BL6 males.

##### RTT + SR (*Mecp2*^+/-^;GFAP-Cre;*Sema3c*^flox/wt^)

RTT + SR mice were generated by first crossing *Mecp2*^+/-^ females to GFAP-Cre males. *Mecp2*^+/-^;Cre+ females were then crossed to *Sema3c*^flox/flox^ males producing four genotypes of offspring: *Mecp2* wildtype, GFAP-Cre negative, *Sema3c*^flox/wt^ (referred to as WT), *Mecp2* wildtype, GFAP-Cre positive, *Sema3c*^flox/wt^ (referred to as WT + SR), *Mecp2* mutant, GFAP-Cre negative, *Sema3c*^flox/wt^ (referred to as RTT), and *Mecp2* mutant, GFAP-Cre positive, *Sema3c*^flox/wt^ (referred to as RTT + SR). Female mice were used to model individuals with RTT, who are predominately female with heterozygous and mosaic MECP2 expression. Male and female mice were tested for visual acuity.

#### Cell culture

##### Primary Astrocyte Culture

Primary astrocyte cultures were isolated using the Miltenyi MACS magnetic cell separation kits. P6-7 wildtype pups were used with 4 animals per prep. Brain cortices were dissected into cold DPBS with meninges and hippocampi removed. Tissue dissociation used the Miltenyi Neural Tissue Dissociation Kit (Miltenyi 130-092-629) and followed manufacturer’s protocol. Cortices were minced and added to a Miltenyi C tube with 1.9 mL of Miltenyi Buffer X, 20 µl Buffer Y, 50 µl enzyme P, and 10 µl enzyme A for papain digestion. Cells were dissociated using the gentleMACS Octo Dissociator (Miltenyi 130-095-937) machine with the provided 37C_NTDK_1 program. After dissociation, the tube was centrifuged for 10 seconds then cells were filtered through the SmartStrainer (Miltenyi 130-098-462) with 10 ml DPBS. Cells were then centrifuged at 300g for 10 minutes and the cell pellet was resuspended in AstroMACS buffer. Astrocytes were isolated using the Miltenyi MACS Anti-ACSA-2 Microbead kit (Miltenyi 130-097-678) following the manufacturer’s protocol with modifications to remove myelin (Miltenyi 130-104-257) and microglia (Miltenyi 130-093-634) through magnetic separation prior to astrocyte positive selection. Cells were counted with Trypan Blue solution (Gibco 15250-061) and plated in a six-well plate prepared the day prior with PDL (Sigma P6407) for 30 minutes, washed 3x with sterile H2O, and dried overnight in the 37-degree incubator. Astrocytes were plated at a density of 40,000 cells per well and grown to confluency over 7-8 days *in vitro* (DIV). Astrocyte base medium contained 50% DMEM (Thermo Fisher Scientific 11960044), 50% Neurobasal (Thermo Fisher Scientific 21103049), Penicillin-Streptomycin (LifeTech 15140-122), Glutamax (LifeTech 35050-061), Sodium Pyruvate (LifeTech 11360-070), N-acetyl-L-cysteine (Sigma A8199), SATO (containing transferrin (Sigma T-1147), BSA (Sigma A-4161), progesterone (Sigma P6149), putrescine (Sigma P5780) and sodium selenite (Sigma S9133)), and heparin binding EGF like growth factor (HbEGF, 5 ng/mL; R&D systems 259-HE/CF). Astrocyte cultures were grown in a humidified incubator at 37C/10% CO_2_ with media changed every 3 days.

##### Collection of astrocyte conditioned media (ACM)

When confluent, astrocytes were washed 3x with warm DPBS to remove excess protein and phenol red. Astrocytes were then grown in astrocyte conditioning media containing 50% DMEM, no phenol red (Thermo Fisher Scientific 21041-025), 50% Neurobasal, no phenol red (Thermo Fisher Scientific 12348-017), Penicillin-Streptomycin (LifeTech 15140-122), Glutamax (LifeTech 35050-061), Sodium Pyruvate (LifeTech 11360-070), N-acetyl-L-cysteine (Sigma A8199), and HbEGF (5 ng/mL; R&D systems 259-HE/CF) for five days. Then conditioned media was collected with a pipette and centrifuged at 3400 RPM for 15 min at 4C to pellet dead cells and debris. Conditioned media was then transferred to a pre-rinsed 3 kDa molecular weight cut-off spin concentrator (Sartorius Vivaspin VS0602) and centrifuged for approximately 90 minutes at 3400 RPM to reduce the volume to 400 microliters. ACM protein concentration was assayed using the Qubit 4 fluorometer (Invitrogen Q33238) with the Qubit protein assay kit (Invitrogen Q33211). ACM was stored up to a week at 4C prior to use.

##### Primary neuron culture

Primary neuron culture used the Miltenyi Neural Tissue Dissociation Kit (Miltenyi 130-092-629) and the Neuron Isolation Kit (130-115-389). Cortices were dissected, digested, and strained as described above for the primary astrocyte isolation. After straining the cells, they were centrifuged for 10 minutes at 300g, supernatant was discarded, and cells were resuspended in 3100 µl of cold DPBS for a debris removal step. 900 µl of Miltenyi Debris Removal solution was added and mixed well. Then, 4 mL of cold DPBS was carefully overlayed to maintain two different phases of solutions. Samples were centrifuged for 10 minutes at 400xg at 4C. The top two phases were aspirated completely then 12 mL of cold DPBS was added. The tube was gently inverted 3x and centrifuged again for 10 minutes at 400xg at 4C. After aspirating supernatant, non-neuronal cell biotin-antibody cocktail and anti-biotin microbeads were used for negative selection of neurons, according to manufacturer’s protocol. Neurons were plated at 40,000 per well in 24 well plates with coverslips (Carolina Biological Supply 633029) coated with PDL (Sigma P6407) and Laminin (Cultrex Trevigen 3400-010-01). Neuron minimal media (MM) contained 50% DMEM, 50% Neurobasal, Penicillin-Streptomycin, Glutamax, sodium pyruvate, N-acetyl-L-cysteine, insulin (Sigma I6634), triiodo-thyronine (Sigma T6397), SATO, B27 (Gibco 17504-044) and forskolin (Sigma F6886). Cortical neuron cultures were maintained in a humidified incubator at 37C/10% CO2.

##### Neurite outgrowth assays

For characterization of SEMA3C impact on outgrowth, primary neurons isolated from wildtype mice were plated on glass coverslips in 24-well plates (three coverslips per condition) with minimal media (MM), MM+ ACM, or MM + ACM + SEMA3C (500ng/ml, R&D 1728-S3-050) for initial characterization of SEMA3C effects on neurite outgrowth. ACM was used at 3 µg/ml concentration. Neurons were cultured for 3 DIV then collected for analysis. Four independent experiments were performed, and analysis of dendrite length was conducted on the average of neurons within an experiment. In each independent experiment, 10 images were taken per condition across three coverslips. The number of neurons analyzed for each replicate was MM: 45, 58, 48, 87; MM + ACM: 48, 73, 41, 107; and MM + ACM + SEMA3C: 53, 48, 26, 93.

For receptor experiments with wildtype ACM, neurons were plated with MM + ACM, MM + ACM + SEMA3C, MM + ACM + SEMA3C + blocking antibody, or MM + ACM + SEMA3C + Normal Goat IgG control (R&D AB-108-C). SEMA3C was used at 200ng/ml and ACM was 3µg/ml. For RTT ACM experiments, wildtype or RTT neurons were plated with MM + RTT ACM, MM + RTT ACM + blocking antibody, or MM + RTT ACM + IgG control. For these assays, RTT neurons were derived from *Mecp2*-null male mice to provide a uniformly MECP2-negative neuronal population. Blocking antibodies were: PLXND1 (R&D AF4160, 5 µg/ml), NRP1 (R&D AF566, 10µg/ml), NRP2 (R&D AF2215, 10 µg/ml), and PLXNA2 (R&D AF5486, 5 µg/ml). IgG control was used at 5 µg/ml for PLXND1 and PLXNA2 experiments and 10 µg/ml for NRP1 and NRP2. Neurons were cultured for 3 DIV then collected for analysis. Four independent experiments were performed for testing each receptor. In each independent experiment, 10 images were taken per condition across three coverslips. The number of neurons analyzed for the four replicates for PLXND1 analysis was ACM: 88, 53, 91, 90; ACM + SEMA3C: 114, 65, 90, 90; ACM + SEMA3C + PLXND1: 80, 73, 92, 60; and ACM + SEMA3C + IgG: 67, 65, 119, 90. For NRP1 analysis, ACM: 71, 51, 99, 72; ACM + SEMA3C: 52, 68, 99, 79; ACM + SEMA3C + NRP1: 64, 68, 84, 35; ACM + SEMA3C + IgG: 52, 68, 95, 47. For NRP2 analysis, ACM: 90, 69, 76, 101; ACM + SEMA3C: 91, 68, 96, 97; ACM + SEMA3C + NRP2: 106, 44, 102, 103; ACM + SEMA3C + IgG: 68, 57, 93, 92.

##### Neurite outgrowth immunohistochemistry and microscopy

Cortical neurons were fixed for 10 minutes with warm 4% paraformaldehyde (PFA) in PBS. Coverslips were washed 3x with room temperature PBS and then blocked with 50% goat serum and 0.5% Triton X-100 in antibody dilution buffer (150 mM NaCl, 50 mM Tris, 100 mM L-lysine, and 1% BSA at pH 7.4) for 30 minutes at room temperature. Coverslips were washed 1x with PBS and incubated overnight at 4C with MAP2 (1:2500, EnCor Biotechnologies CPCA-MAP2) in antibody buffer with 10% goat serum. The next day, cells were washed 3x with PBS and incubated for two hours with goat anti-chicken Alexa-Fluor 488 (1:1000, Thermo Fisher Scientific A11039) in antibody buffer with 10% goat serum. Coverslips were washed 3x with PBS and mounted with SlowFade + DAPI on glass slides sealed with nail polish.

Imaging was performed with a Plan-Apochromat 10x/0.45 M27 objective on an Axio Imager 7.2 fluorescent microscope and AxioCam HR3 (Zeiss) with at least one neuron per image. Quantification of dendrite length was conducted in FIJI using the NeuronJ plugin. Four independent experiments (biological replicates) were conducted for each condition and statistical analysis of dendrite length was performed on the average of cells within a biological replicate.

#### Electrophysiology

##### Acute slice preparation

Coronal brain slices were prepared from P60 WT, WT + SR, RTT, and RTT + SR mice. Animals were deeply anesthetized by injection with 0.4g/kg Avertin solution and decapitated. The brain was removed and cut into 300 µm coronal sections using a Leica VT1000s vibratome. The brain dissection was performed at physiological temperature (37C), with choline chloride-based dissection solution composed of (in mM): 93 choline chloride, 2.5 KCl, 2 MgCl_2_, 2 CaCl_2_, 1.2 NaH_2_PO_4_, 20 HEPES, 5 Sodium L-ascorbate, 25 D-glucose and 24 NaHCO3. Slices were then placed in a recovery chamber containing HEPES artificial cerebrospinal fluid (aCSF) composed of (in mM): 5 HEPES, 93 NaCl, 2.5 KCl, 2 MgCl_2_, 2 CaCl_2_, 1.2 NaH_2_PO_4_, 5 Sodium L-ascorbate, 25 D-glucose and 24 NaHCO3. Slices recovered for 10 min at 37C and then for at least 50 min at room temperature before recordings were performed for 4-6 hours after slicing. Solutions were supplemented with 1 mM sodium pyruvate and continuously equilibrated with carbogen (95% O2/ 5% CO2).

##### Electrophysiology recordings

Slices were placed in a recording chamber and perfused with carbogen-saturated aCSF composed of (in mM): 126 NaCl, 2.5 KCl, 1.2 MgCl_2_, 2.5 CaCl_2_, 1.2 NaH_2_PO_4_, 11 D-glucose and 25 NaHCO3, supplemented with 1 mM sodium pyruvate. Whole-cell patch-clamp recordings were performed in the soma of visually identified pyramidal neurons from L2/3 of the visual cortex using infrared camera on a Scientifica microscope. Patch pipettes (borosilicate glass pipette; Harvard Apparatus #BS4 64-0805; 4-6 MΩ of resistance) were filled with an internal solution composed of (in mM): 105 K-gluconate, 30 KCl, 10 phosphocreatine, 10 HEPES, 4 ATP-Mg, 0.3 GTP-Tris, 0.3 EGTA and 0.2% biocytin, adjusted to pH 7.2 with KOH. Recordings were performed using an Axon Digidata 1550A and a Multiclamp 700B amplifier (Molecular Devices). All recordings were sampled at 10 kHz. Spontaneous excitatory postsynaptic currents (sEPSCs) were recorded at a membrane holding potential of –60 mV, and spontaneous inhibitory postsynaptic currents (sIPSCs) were recorded at a membrane holding potential of 0 mV. All currents were recorded at room temperature (22–24C), and only one neuron was patched in each slice. Once the whole-cell configuration was attained, we waited 5 minutes for the baseline to stabilize before recording sEPSCs for 5 minutes, then after switching the holding potential, we waited another 5 minutes before recording sIPSCs for 5 minutes as well. Recordings were discarded if the access resistances were > 20 MΩ or changed > 25% during the recording. After recording, the pipette was carefully removed from the slices containing a biocytin-filled neuron, and the slices were transferred to PBS with 4% PFA. Electrophysiology recording analyses were performed using MiniAnalysis version 6.0.8 (Synaptosoft) and currents were detected after using a 1 kHz post-hoc filter. 1-4 neurons were recorded from each animal with animal averages used for statistics.

#### *Ex vivo* dendrite and spine analysis

Following electrophysiology experiments, the brain slices were post-fixed in 4% PFA in PBS overnight at 4 degrees. The following day, the sections were washed with PBS 3x and stored for up to one week in PBS at 4 degrees. For immunostaining, sections were blocked for 3 hours at RT with 10% Normal goat serum and 0.5% Triton-X 100 in PBS. Sections were incubated with primary antibodies overnight at room temperature in blocking buffer with 1:1000 Rabbit anti-MECP2 (Cell Signaling #3456) and 1:1000 mouse anti-S100B (Sigma #2532) to label astrocytes. The following day, sections were washed 3x 10 minutes with PBS followed by 4 hours in secondary in buffer consisting of 5% NGS and 0.2% Triton-X 100 with Alexa Goat anti-Rabbit 488 (1:1000), Alexa Goat anti-Mouse 647 (1:1000), and conjugated Streptavidin-Alexa 594 (1:1500). Sections were washed 3x 10 minutes in PBS and then incubated with DAPI (1:10,000 in PBS) before mounting with Fluoromount mounting media.

##### Dendrite arborization

Neurons were imaged with a Plan-Apochromat 10x/0.45 M27 objective on an Axio Imager 7.2 fluorescent microscope and AxioCam HR3 (Zeiss) with one image encompassing the soma and basal dendrites and one image encompassing the apical dendrites and edge of the soma. Z-stacks of 0.5 micron increments were taken with the number of steps required to image the entire soma and dendrites. All neurons were confirmed to be in L2/3 by DAPI staining. Sholl analysis was conducted in FIJI by counting the number of crossings of concentric circles at 10-micron intervals from the soma. Statistics were conducted with each animal as N (average of 1-4 neurons per animal) using mixed-effects models (restricted maximum likelihood estimation) in GraphPad Prism version 10.5.0. This approach was used, rather than a traditional two-way ANOVA because some neurons did not extend dendrites to the most distal radii, resulting in missing values at greater distances from the soma across genotypes. The mixed-effects model with REML accommodates these missing observations while testing for main effects and interactions. Post-hoc pairwise comparisons were adjusted for multiple testing using Tukey’s correction.

##### Spine density and morphology

Dendrites were imaged with a Plan-Apochromat 63x/1.4 Oil DIC M27 objective on a Zeiss 880 AxioObserver with Airyscan detector. For basal dendrites, only primary dendrites were imaged. For apical dendrites, only dendrites that reached the apical surface of the brain were imaged. Five basal and apical dendrites were imaged for each neuron. Z-stacks of 0.20 micron increments were taken with the number of steps required to encompass the entire dendrite and spines. Laser settings and gain were consistent throughout imaging. Images were Airyscan processed in Zen and exported as TIFF files. Spine analysis and morphology were analyzed using NeuronStudio. Spines were automatically labeled by NeuronStudio and verified by a trained experimenter, blinded to genotype. Spine density was quantified as the number of spines per micron and analysis was performed on the average of basal or apical dendrites per mouse. Spine morphology was based on classifications previously described^105^.

##### Soma size, nucleus size, and quantification of MECP2

For soma and nucleus size analysis, neurons were imaged with a Plan-Apochromat 40x/1.4 Oil DIC M27 objective on a Zeiss 900 Axio Observer.Z1 with the soma centered within the image. Z-stacks of 0.5 micron increments were taken with 15 steps per image. Soma and nucleus size were measured on a maximum intensity projection in FIJI/ImageJ. Blinded to genotype, the experimenter outlined each soma or nucleus three times, recorded the area, and averaged for each neuron. These images were then used for MECP2 quantification using the CellCounter plugin in FIJI/ImageJ.

#### Single molecule fluorescent in situ hybridization (smFISH)

Mice were anesthetized with an intraperitoneal injection of 100mg/kg ketamine (Victor Medical Company 1699053) and 20mg/kg xylazine (Victor Medical Company 1264078) and perfused transcardially with 4% PFA in PBS. Brains were extracted, postfixed in 4% PFA (Electron Microscopy Sciences 19210) overnight at 4C, cryoprotected in 30% sucrose in PBS for 48-72 hours, and embedded in OCT compound (Tissue-Tek). Coronal brain sections were cryosectioned at 16 μm onto slides (Superfrost Plus, catalog #48311-703, VWR). Slides were warmed for 30 minutes on a slide warmer set at 60°C and then processed using RNAscope Multiplex Fluorescent assay kit (ACD Biotechne 323100) following manufacturer’s protocol. *Sema3c* (441441) and GLAST (*Slc1a3*, 430781) probes were used. For analysis, layer 2/3 of the visual cortex was imaged with a Plan-Apochromat 40x/1.4 Oil DIC M27 objective on a Zeiss 900 Axio Observer.Z1. Z-stacks of 0.5 micron increments were taken with 8 steps per image. Blinded to genotype, *Slc1a3+* astrocyte nuclei (labeled with DAPI) were first identified using the CellCounter plugin in FIJI/ImageJ. Then the number of *Sema3c* puncta per astrocyte nucleus was recorded. Four images were taken per animal and averaged for the animal mean used for statistical analysis.

#### Cortical width

Cortical width was assessed on adult brain tissue from mice used for behavioral analysis. Samples were prepared as for single molecule fluorescent in situ hybridization. Slides were immunostained with 1:1000 Rabbit anti-MECP2 (Cell Signaling #3456) and DAPI. Slides were imaged with a Plan-Apochromat 10x/0.45 M27 objective on an Axio Imager 7.2 fluorescent microscope and AxioCam HR3 (Zeiss) with the visual cortex tiled at 10% overlap. Blinded to genotype, the experimenter measured the width of layer 1 and L2/3 three times per image, recorded the length in microns, and then averaged for each image. Three images were taken per animal and averaged for the animal mean used for statistical analysis.

#### Mouse behavior

All mouse behavior experiments were approved by Salk IACUC and conducted in the Salk behavior core, In Vivo Scientific Services. Behavior was conducted during the light cycle with optomotor performed between Zeitgeber time 2-6 and open field and rotarod conducted between Zeitgeber time 8-11. All assays were run and analyzed by a trained experimenter blinded to genotype. Elevated plus maze was conducted at P45. Optomotor assay, open field, and rotarod were run between P120 and P125 in that order, on different days. Visual acuity in male mice was conducted at P45.

##### Elevated plus maze

The mouse was placed in the center of a ‘plus’-shaped apparatus (approximately 70 cm wide x 116 cm tall) with two ‘open’ arms with no walls and two ‘closed’ arms with walls. A camera above the apparatus and ANY-Maze software (Stoelting Co.) is used to track and quantify the mouse’s activity and location over a five-minute period.

##### Optomotor assay

The optomotor assay to measure visual acuity used the OptoMotry VR apparatus (CerebralMechanics Inc.), which consists of an arena of four computer monitors with a pedestal for the mouse in the center. A video camera is positioned directly above the animal. The entire apparatus is inside a sound-proof chamber. A virtual cylinder with vertical sine wave grating is projected in two-dimensional coordinate space on the computer monitors providing the visual stimulus that the mouse tracks with a reflexive head movement^110^. A low spatial frequency (0.42 cycle/degree) sine wave grating at 100% contrast is projected and a trained observer assesses “yes” or “no” for tracking behavior within a few seconds. Stimuli are then presented using the staircase method with increasing spatial frequencies presented (from 0.42 to 0.042 cycles per degree) until the mouse no longer responds. The threshold for clockwise and counterclockwise stimuli is determined and then averaged for the mean threshold.

##### Open field

The open field assay consists of a square arena (43.2 × 43.2 × 30.5 cm; Med Associates Inc. ENV-515S-A) in which the mouse’s activity is monitored. On the day of the assay, mice were habituated to the testing room for 30 minutes in their home cage. The room lighting was set to approximately 30 lux. The mouse was then placed into the center of the arena and activity was automatically tracked by Activity Monitor software (Med Associates) for 60 minutes. Distance traveled (horizontal beam breaks), rearing (vertical beam breaks), and percentage of time spent in the center were quantified.

##### Rotarod

The Rotarod apparatus (SD Instruments) consists of a rotating rod set to an accelerating speed. On day 1, mice are habituated to the rod at a rotating speed of 4 RPM. During this training, mice are exposed to the rod for 5 minutes. Mice that fall during the training are placed back on the rod until the training session time has elapsed, or the mouse has stayed on the rod for two consecutive minutes. The following day, mice are placed back on the rod during which the speed increases from 4 to 30 RPM over 5 minutes. Mice are tested three times with a 5 minute rest period in their home cage in between trials. The average latency to fall and total distance traveled across three trials is reported.

### Statistical analysis

Statistical analyses were performed in GraphPad Prism (version 10.5.0). Significance was set at alpha = 0.05. For comparison of two groups, T-test was used for normally distributed data. For comparison of three or more groups for cell culture experiments, one-way ANOVA with Tukey’s multiple comparisons. For comparison of four groups for *ex vivo* and *in vivo* studies, two-way ANOVA was used to test for *Mecp2* and SEMA3C effects, alongside Tukey’s multiple comparisons for pairwise comparison. For data where animal averages comprised different numbers of data points (dendrite arborization, spine analysis and electrophysiology), nested ANOVAs were also calculated and are included in the statistics table (Supplementary Table 1). Spine morphology was analyzed with by three-way ANOVA (factors: spine type, *Mecp2,* SEMA3C). Distributions for electrophysiology data were analyzed used pairwise Kolmogorov–Smirnov tests. Graphs were generated in Prism.

**Supplementary data table 1.** Statistical tests and summary statistics from studies.

---

**Figure S1.**
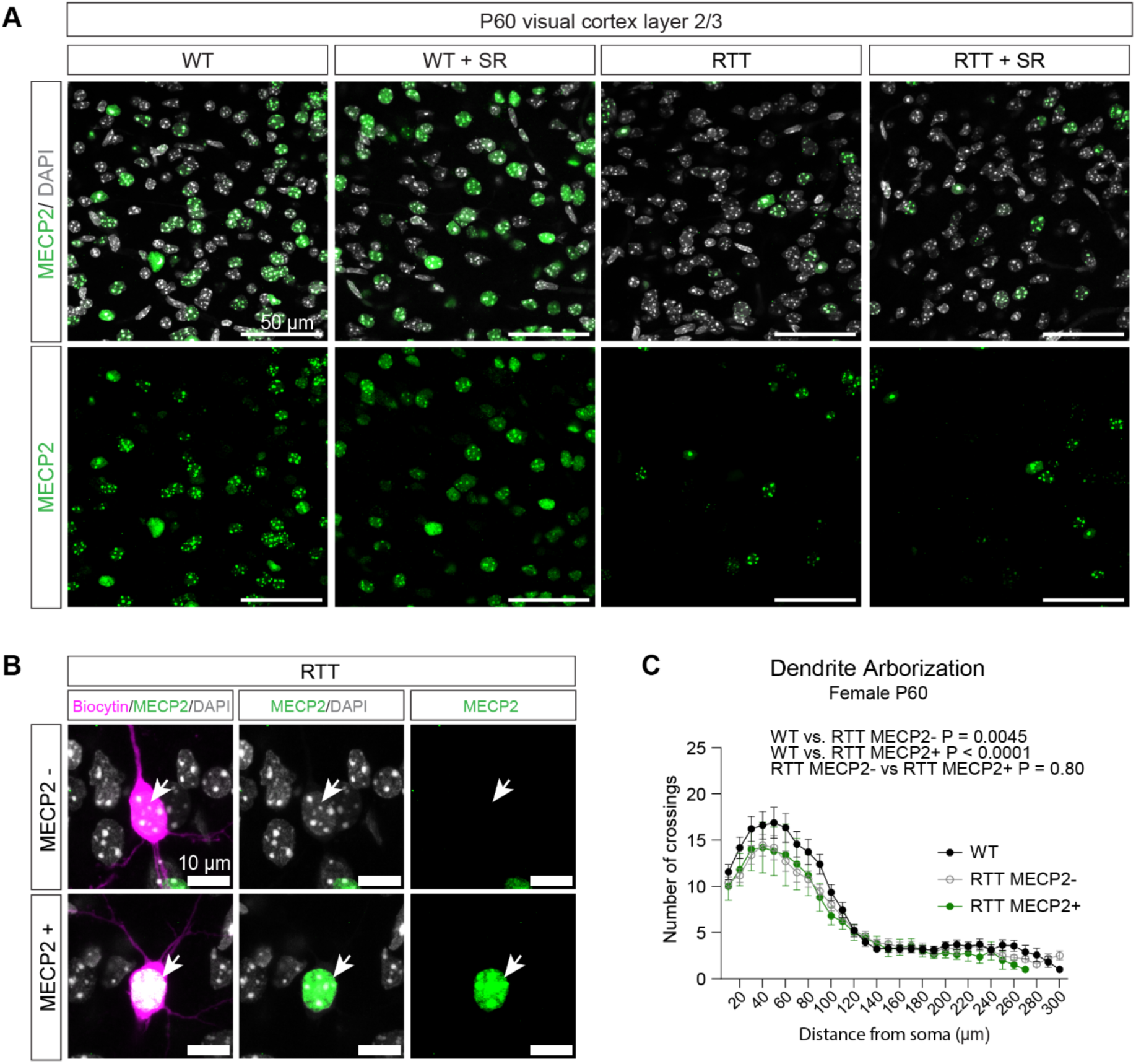
MECP2 staining and dendrite arborization, related to Figure 2. **(A)** Representative images of immunostaining for MECP2 (green) and DAPI (gray) in VC L2/3 at P60. Scale bar 50 µm. **(B)** Representative images of MECP2 negative (top) and MECP2 positive (bottom) P60 pyramidal neuron soma (biocytin, magenta) with immunostaining for MECP2 (green) and DAPI (gray), scale bar is 10 µm. Arrowhead denotes labeled neuron soma. **(C)** Quantification of dendrite arborization in WT, RTT MECP2 negative (-), and RTT MECP2 positive (+) neurons. N = 18 WT, 18 RTT MECP2-, and 5 RTT MECP2+. Mixed-effects model (REML), two factors: genotype and distance from soma; Tukey’s multiple comparisons. P values denoted on figure.

**Figure S2.**
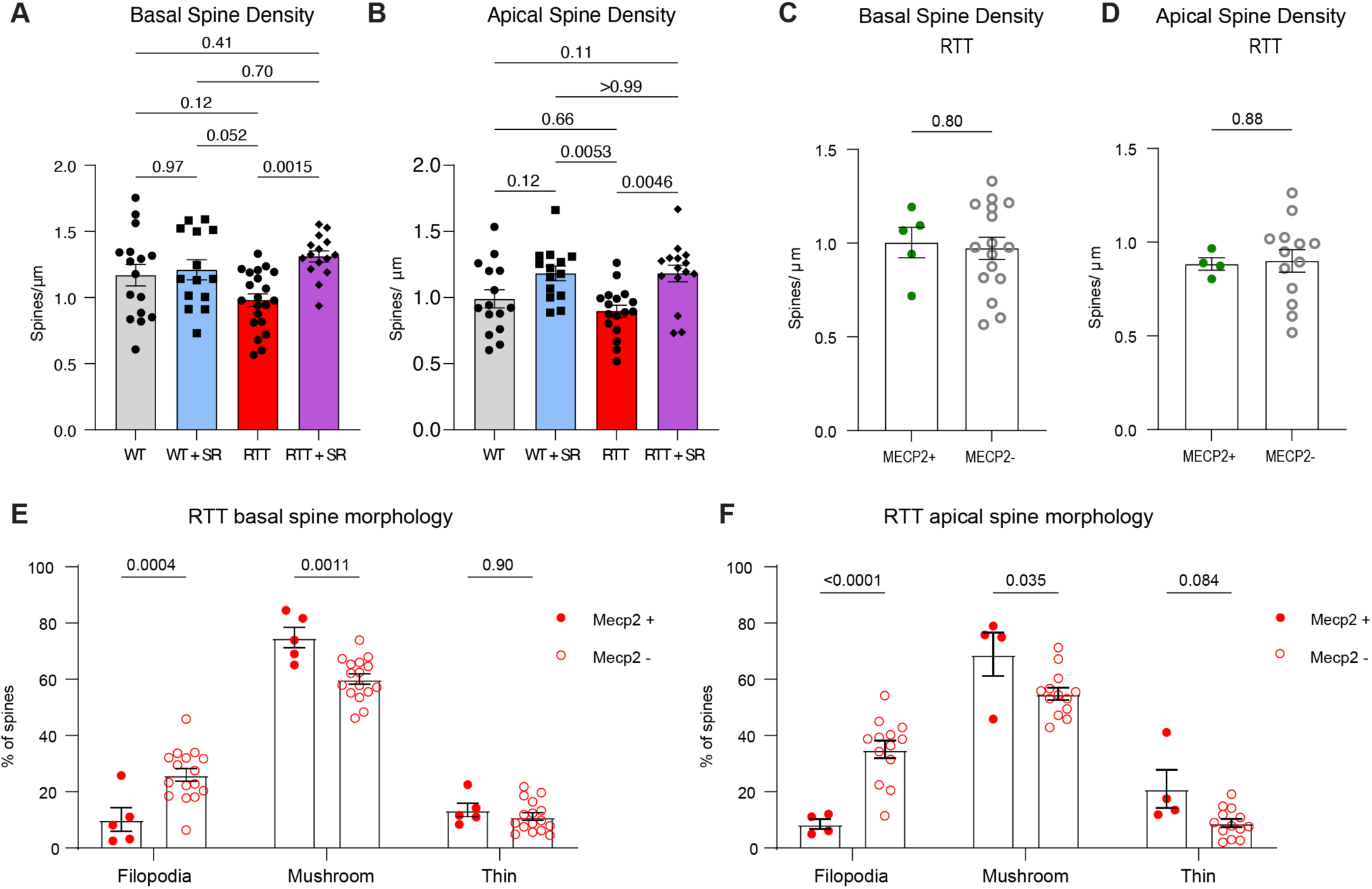
Additional characterization of spine density and spine morphology, related to Figure 3. **(A-B)** Quantification of spine density on a per neuron basis. Each symbol represents one neuron. For basal (A), N = 16 WT, 14 WT + SR, 21 RTT, and 15 RTT + SR neurons. For apical (B), N = 15 WT, 14 WT + SR, 17 RTT, and 15 RTT + SR neurons. Two-way ANOVA with Tukey’s multiple comparisons. **(C-D)** Comparison of spine density in MECP2+ versus MECP2- neurons in RTT mice. For basal (C), N = 5 MECP2+ neurons and 16 MECP2- neurons. For apical (D), N = 4 MECP2+ neurons and 13 MECP2- neurons. Unpaired T-test. **(E-F**) Comparison of spine morphology in MECP2+ versus MECP2- neurons in RTT mice for filopodia, mushroom, and thin spine types. For basal (E), N = 5 MECP2+ neurons and 16 MECP2- neurons. For apical (F), N = 4 MECP2+ neurons and 13 MECP2- neurons. Each symbol represents one neuron. Two-way ANOVA with Sidak’s multiple comparisons. A-F. P values denoted. Error bars depict SEM.

**Figure S3.**
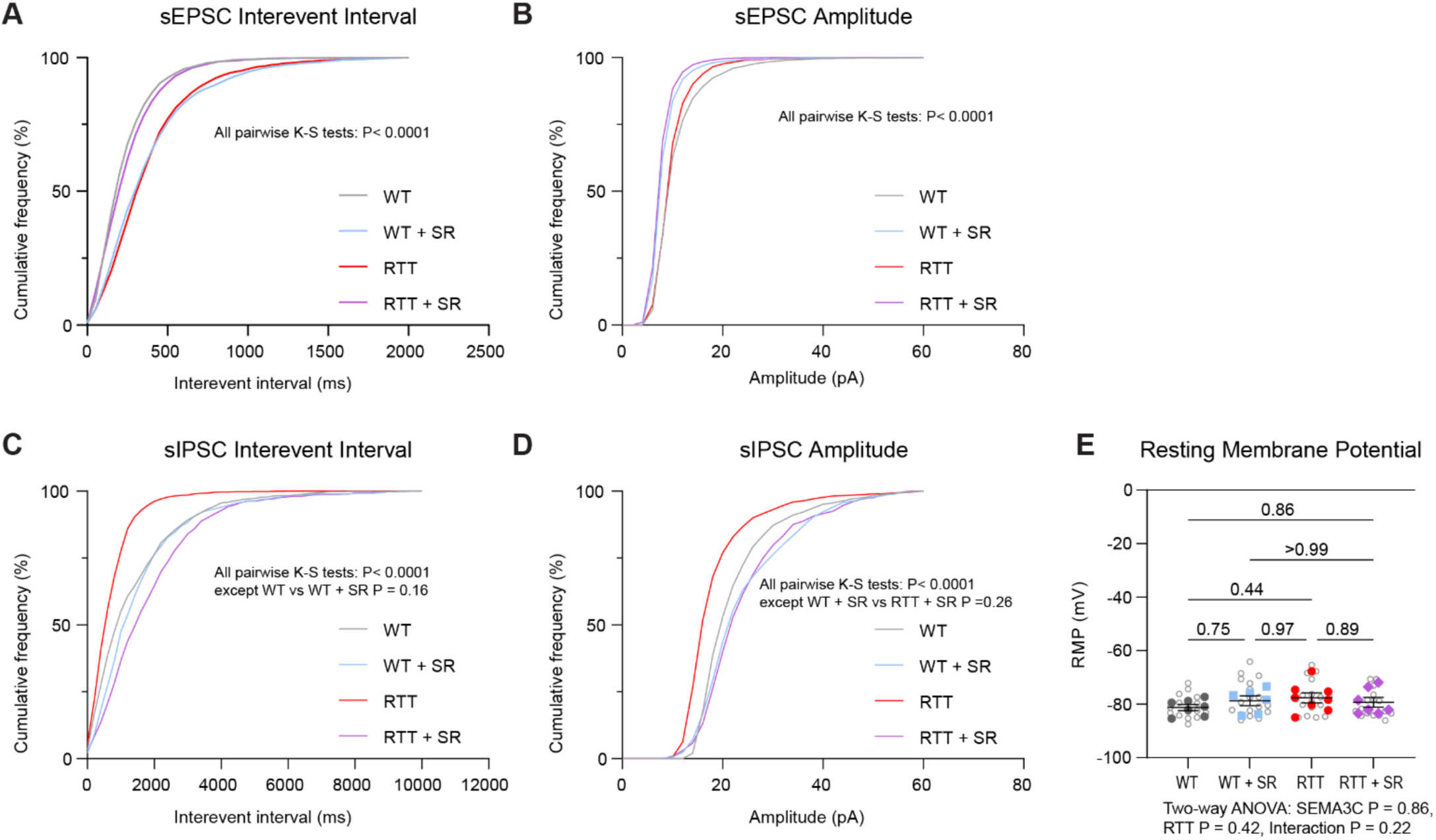
Cumulative frequency distributions and resting membrane potential, related to Figure 4. **(A)** Cumulative frequency distribution of sEPSC interevent intervals (inverse of frequency) for all events. **(B)** Cumulative frequency distribution of sEPSC amplitude for all events. **(C)** Cumulative frequency distribution of sIPSC interevent intervals for all events. **(D)** Cumulative frequency distribution of sIPSC amplitude for all events. A-D. Kolmogorov-Smirnov tests. (**E)** Average resting membrane potential (mV). Gray open circles represent individual cells. Solid dots represent animal averages. N = 7 WT, 6 WT + SR, 8 RTT, and 7 RTT + SR animals with 1-4 cells per animal. Two-way ANOVA on per-mouse means with Tukey’s multiple comparisons. P values denoted. Error bars depict mean ± SEM.

**Figure S4.**
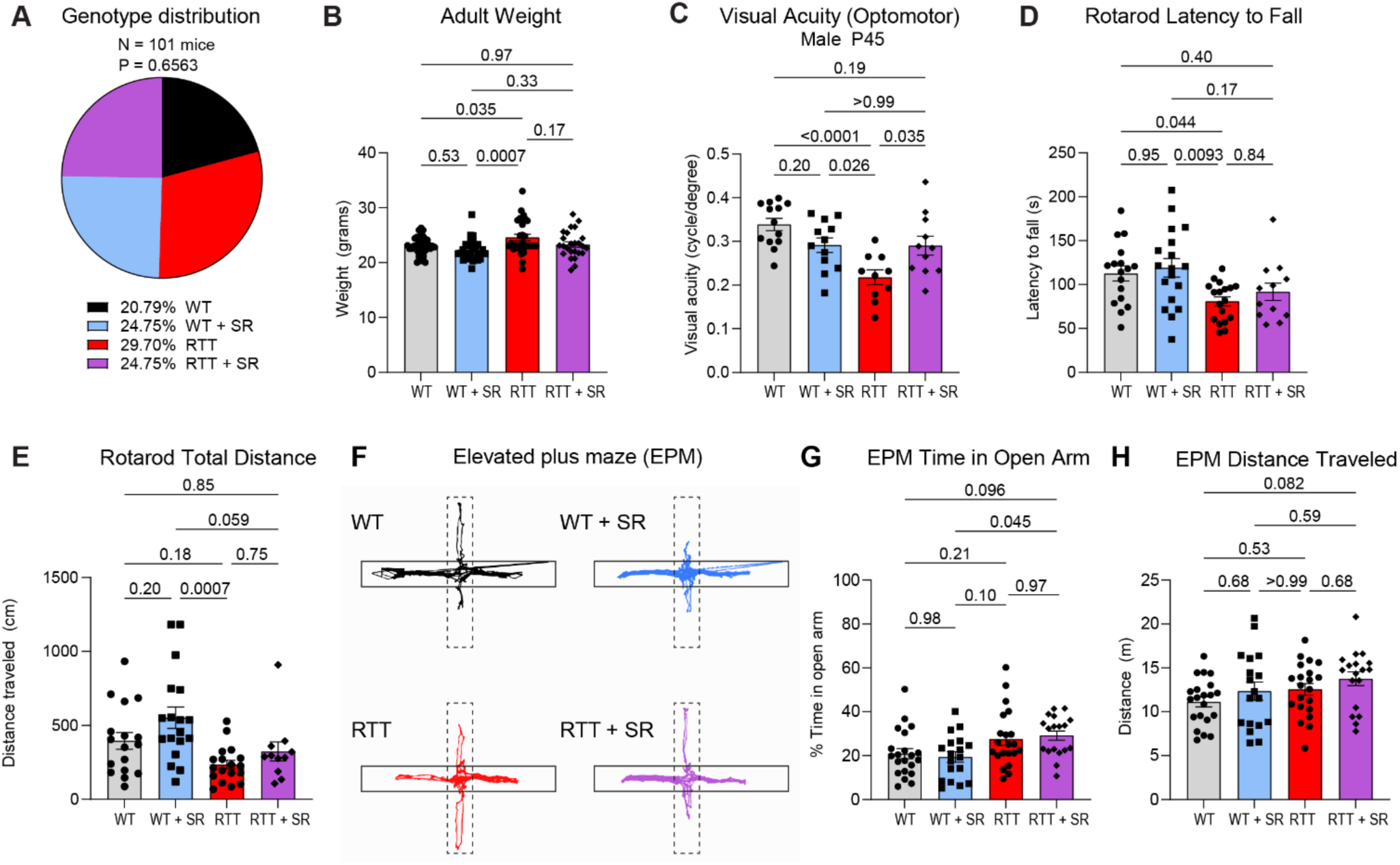
Additional characterization of RTT + SR behavior, related to Figure 5. **(A)** Distribution of genotypes. Chi-square test for observed distribution versus expected distribution (25% per genotype). P value denoted. **(C)** Adult (P120) female body weight. N = 35 WT, 32 WT + SR, 29 RTT, and 25 RTT + SR **C.** Visual acuity threshold (cycles/degree) in male *Mecp2* null mice at P45. N = 13 WT, 12 WT + SR, 10 RTT, and 11 RTT + SR. **D.** Average latency to fall from the rotarod. **E.** Total distance traveled on the rotarod. **D-E.** N = 17 WT, 18 WT + SR, 18 RTT, and 12 RTT + SR females at P120-125. **F.** Representative traces of elevated plus maze activity (EPM), dotted lines indicate open arms while closed boxes indicate closed arms. **G.** Percentage of EPM assay time spent in open arms. **H.** Total distance traveled during the EPM assay. **G-H.** N = 15 WT, 14 WT + SR, 15 RTT, 12 RTT + SR females at P45. **B-E, G-H.** Two-way ANOVA with Tukey’s multiple comparisons. P values denoted. Error bars depict SEM. Each dot represents one animal.

**Figure S5.**
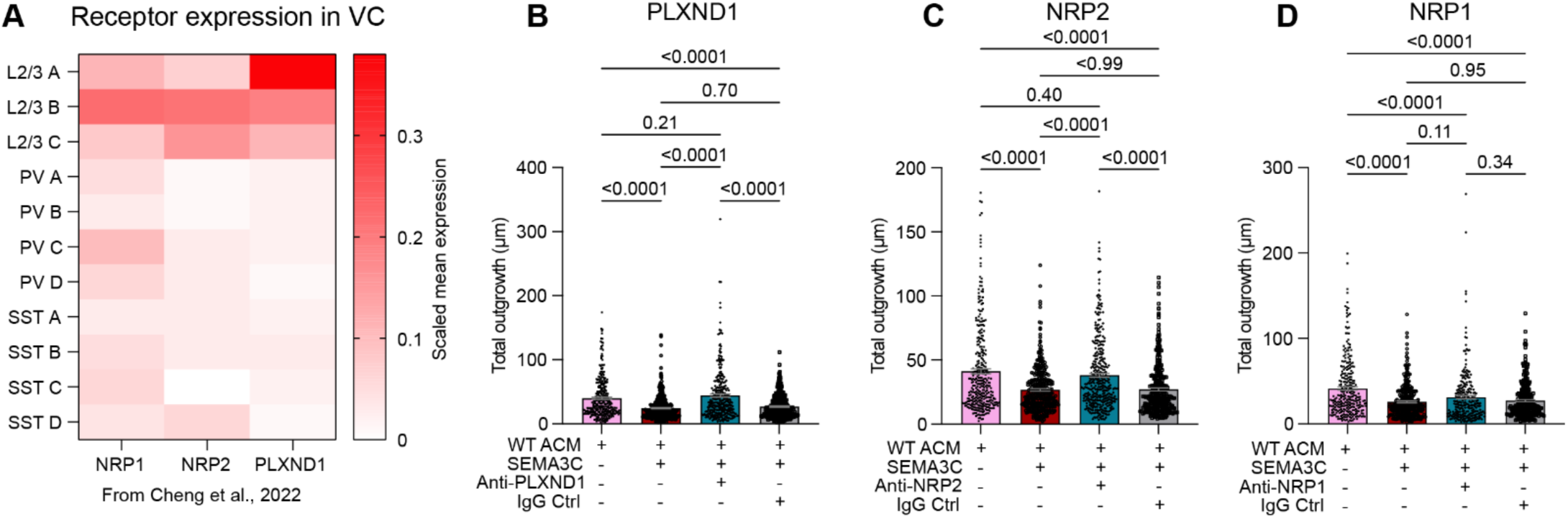
Receptor expression profile in cortical neurons and dendrite outgrowth for individual neurons, related to Figure 6. **(A)** NRP1, NRP2, and PLXND1 expression in VC neuron types from Cheng et al., 2022. L2/3 = L2/3 excitatory neurons and PV and SST = inhibitory neuron subtypes. **B-D.** Quantification of dendrite outgrowth on a per neuron basis for receptor blocking experiments, across 4 independent experiments with 3 coverslips per condition: **(B)** PLXND1. N = 322 neurons for ACM, 359 for ACM + SEMA3C, 305 for ACM + SEMA3C + PLXND1, and 341 for ACM + SEMA3C + IgG conditions. **(C)** NRP2. N = 336 neurons for ACM, 352 for ACM + SEMA3C, 355 for ACM + SEMA3C + NRP2, 310 for ACM + SEMA3C + IgG conditions. **(D)** NRP1. N = 293 for ACM, 298 for ACM + SEMA3C, 251 for ACM + SEMA3C + NRP1, and 262 for ACM + SEMA3C + IgG. **B-D:** One-way ANOVA with Tukey’s multiple comparisons. P values denoted. Error bars show SEM. Each dot represents one neuron.

**S6.**
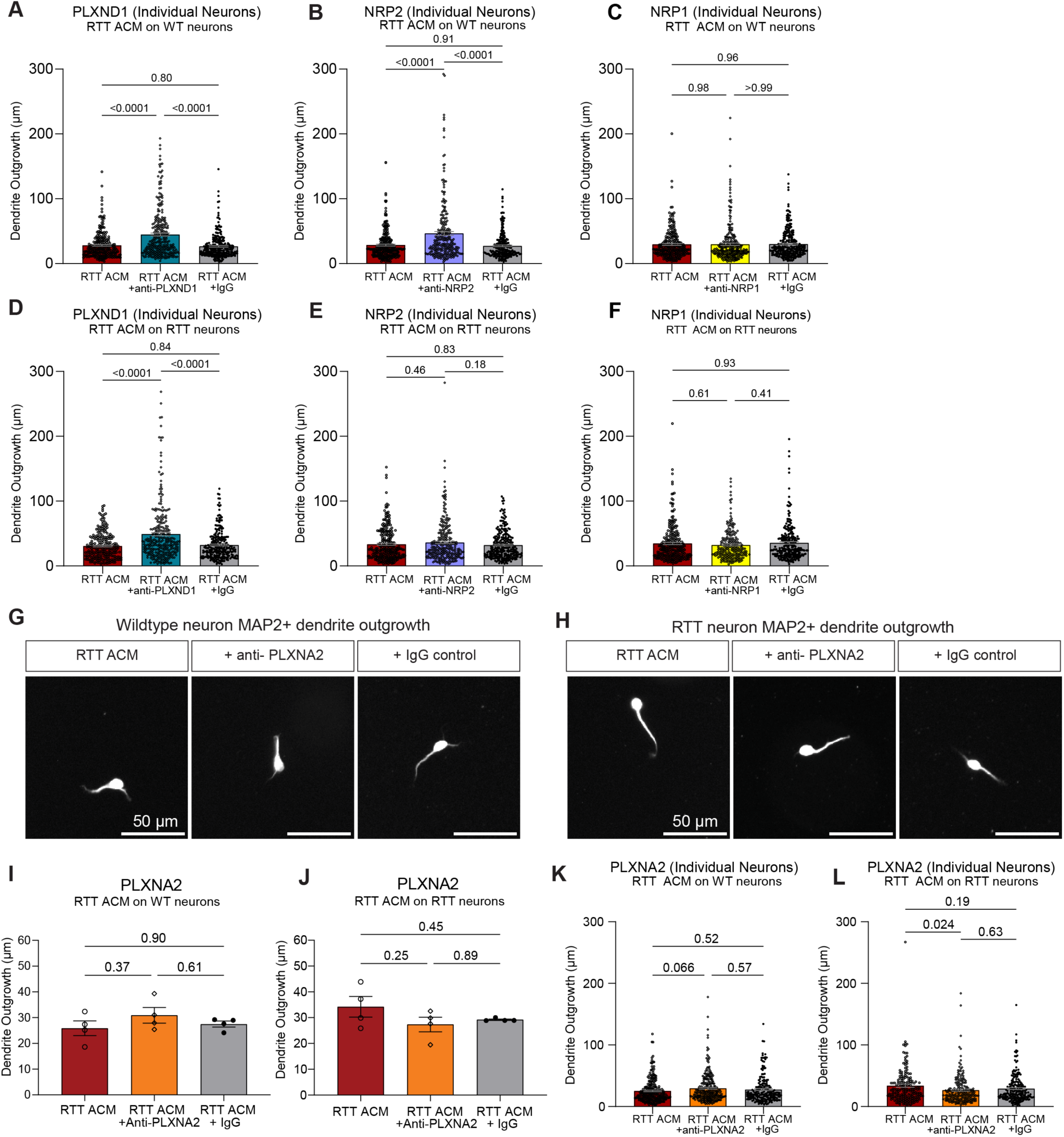
Dendrite outgrowth for individual neurons and blocking PLXNA2 in RTT ACM does not improve dendrite outgrowth, related to Figure 7. **A-F.** Quantification of dendrite outgrowth on a per neuron basis for receptor blocking experiments with RTT ACM on WT or RTT neurons, across 4 independent experiments with 3 coverslips per condition: **(A)** PLXND1 in RTT ACM on WT neurons. N = 242 neurons for RTT ACM, 287 for RTT ACM + PLXND1, 260 for RTT ACM + IgG, **(B)** NRP2 in RTT ACM on WT neurons. N = 263 neurons for RTT ACM, 258 RTT ACM + NRP2, 220 for RTT ACM + IgG, **(C)** NRP1 in RTT ACM on WT neurons. N = 242 neurons for RTT ACM, 271 RTT ACM + NRP1, 267 RTT ACM +IgG, **(D)** PLXND1 in RTT ACM on RTT neurons. N = 238 neurons for RTT ACM, 280 RTT ACM + PLXND1, 247 RTT ACM + IgG, **(E)** NRP2 in RTT ACM on RTT neurons. N = 263 neurons for RTT ACM, 267 RTT ACM + NRP2, 261 ACM + IgG, **(F)** NRP1 in RTT ACM on RTT neurons. N = 264 neurons for RTT ACM, 219 RTT ACM + NRP2, 247 RTT ACM + IgG. **(G-H)** Representative images of (G) WT neuron dendrite outgrowth or (H) RTT neuron dendrite outgrowth with RTT ACM (left panels), RTT ACM with blocking antibody to PLXNA2 (middle panels, 5µg/ml), and IgG control (right panels, 5µg/ml). Scale bars are 50 µm. **(I-J)** Quantification of average total dendrite outgrowth for RTT ACM alone, with blocking antibody for PLXNA2 or with IgG control on (I) WT neurons or (J) RTT neurons. **(K-L)** Quantification of dendrite outgrowth on a per neuron basis for receptor blocking experiments with RTT ACM on WT or RTT neurons, across 4 independent experiments with 3 coverslips per condition: (K) PLXNA2 in RTT ACM on WT neurons. N = 237 neurons for RTT ACM, 274 for RTT ACM + anti-PLXNA2, 201 for RTT ACM + IgG, and (L) PLXNA2 in RTT ACM on RTT neurons. N = 190 neurons for RTT ACM, 215 for RTT ACM + anti-PLXNA2, 215 for RTT ACM + IgG. A-F, I-L: One-way ANOVA with Tukey’s multiple comparisons. P values denoted. Error bars show SEM. Each dot represents one neuron.

## References

1. Amir, R.E., Van den Veyver, I.B., Wan, M., Tran, C.Q., Francke, U., and Zoghbi, H.Y. (1999). Rett syndrome is caused by mutations in X-linked MECP2, encoding methyl-CpG-binding protein 2. Nat Genet 23, 185–188. 10.1038/13810.

2. Chahrour, M., and Zoghbi, H.Y. (2007). The story of Rett syndrome: from clinic to neurobiology. Neuron 56, 422–437. 10.1016/j.neuron.2007.10.001.

3. Sugino, K., Hempel, C.M., Okaty, B.W., Arnson, H.A., Kato, S., Dani, V.S., and Nelson, S.B. (2014). Cell-type-specific repression by methyl-CpG-binding protein 2 is biased toward long genes. J Neurosci 34, 12877–12883. 10.1523/JNEUROSCI.2674-14.2014.

4. Chahrour, M., Jung, S.Y., Shaw, C., Zhou, X., Wong, S.T., Qin, J., and Zoghbi, H.Y. (2008). MeCP2, a key contributor to neurological disease, activates and represses transcription. Science 320, 1224–1229. 10.1126/science.1153252.

5. Bach, S., Ryan, N.M., Guasoni, P., Corvin, A.P., El-Nemr, R.A., Khan, D., Sanfeliu, A., and Tropea, D. (2020). Methyl-CpG-binding protein 2 mediates overlapping mechanisms across brain disorders. Sci Rep 10, 22255. 10.1038/s41598-020-79268-0.

6. Urdinguio, R.G., Lopez-Serra, L., Lopez-Nieva, P., Alaminos, M., Diaz-Uriarte, R., Fernandez, A.F., and Esteller, M. (2008). Mecp2-null mice provide new neuronal targets for Rett syndrome. PLoS One 3, e3669. 10.1371/journal.pone.0003669.

7. Yasui, D.H., Xu, H., Dunaway, K.W., Lasalle, J.M., Jin, L.W., and Maezawa, I. (2013). MeCP2 modulates gene expression pathways in astrocytes. Mol Autism 4, 3. 10.1186/2040-2392-4-3.

8. Colantuoni, C., Jeon, O.H., Hyder, K., Chenchik, A., Khimani, A.H., Narayanan, V., Hoffman, E.P., Kaufmann, W.E., Naidu, S., and Pevsner, J. (2001). Gene expression profiling in postmortem Rett Syndrome brain: differential gene expression and patient classification. Neurobiol Dis 8, 847–865. 10.1006/nbdi.2001.0428.

9. Forbes-Lorman, R.M., Kurian, J.R., and Auger, A.P. (2014). MeCP2 regulates GFAP expression within the developing brain. Brain Res 1543, 151–158. 10.1016/j.brainres.2013.11.011.

10. Pacheco, N.L., Heaven, M.R., Holt, L.M., Crossman, D.K., Boggio, K.J., Shaffer, S.A., Flint, D.L., and Olsen, M.L. (2017). RNA sequencing and proteomics approaches reveal novel deficits in the cortex of Mecp2-deficient mice, a model for Rett syndrome. Mol Autism 8, 56. 10.1186/s13229-017-0174-4.

11. Delepine, C., Nectoux, J., Letourneur, F., Baud, V., Chelly, J., Billuart, P., and Bienvenu, T. (2015). Astrocyte Transcriptome from the Mecp2(308)-Truncated Mouse Model of Rett Syndrome. Neuromolecular Med 17, 353–363. 10.1007/s12017-015-8363-9.

12. Caldwell, A.L.M., Sancho, L., Deng, J., Bosworth, A., Miglietta, A., Diedrich, J.K., Shokhirev, M.N., and Allen, N.J. (2022). Aberrant astrocyte protein secretion contributes to altered neuronal development in multiple models of neurodevelopmental disorders. Nat Neurosci 25, 1163–1178. 10.1038/s41593-022-01150-1.

13. Okabe, Y., Takahashi, T., Mitsumasu, C., Kosai, K., Tanaka, E., and Matsuishi, T. (2012). Alterations of gene expression and glutamate clearance in astrocytes derived from an MeCP2-null mouse model of Rett syndrome. PLoS One 7, e35354. 10.1371/journal.pone.0035354.

14. Kahanovitch, U., Cuddapah, V.A., Pacheco, N.L., Holt, L.M., Mulkey, D.K., Percy, A.K., and Olsen, M.L. (2018). MeCP2 Deficiency Leads to Loss of Glial Kir4.1. eNeuro 5. 10.1523/ENEURO.0194-17.2018.

15. Ballas, N., Lioy, D.T., Grunseich, C., and Mandel, G. (2009). Non-cell autonomous influence of MeCP2-deficient glia on neuronal dendritic morphology. Nat Neurosci 12, 311–317. 10.1038/nn.2275.

16. Lioy, D.T., Garg, S.K., Monaghan, C.E., Raber, J., Foust, K.D., Kaspar, B.K., Hirrlinger, P.G., Kirchhoff, F., Bissonnette, J.M., Ballas, N., and Mandel, G. (2011). A role for glia in the progression of Rett’s syndrome. Nature 475, 497–500. 10.1038/nature10214.

17. Maezawa, I., Swanberg, S., Harvey, D., LaSalle, J.M., and Jin, L.W. (2009). Rett syndrome astrocytes are abnormal and spread MeCP2 deficiency through gap junctions. J Neurosci 29, 5051–5061. 10.1523/JNEUROSCI.0324-09.2009.

18. Sun, J., Osenberg, S., Irwin, A., Ma, L.H., Lee, N., Xiang, Y., Li, F., Wan, Y.W., Park, I.H., Maletic-Savatic, M., and Ballas, N. (2023). Mutations in the transcriptional regulator MeCP2 severely impact key cellular and molecular signatures of human astrocytes during maturation. Cell Rep 42, 111942. 10.1016/j.celrep.2022.111942.

19. Chung, W.S., Baldwin, K.T., and Allen, N.J. (2024). Astrocyte Regulation of Synapse Formation, Maturation, and Elimination. Cold Spring Harb Perspect Biol 16. 10.1101/cshperspect.a041352.

20. Allen, M., Huang, B.S., Notaras, M.J., Lodhi, A., Barrio-Alonso, E., Lituma, P.J., Wolujewicz, P., Witztum, J., Longo, F., Chen, M., et al. (2022). Astrocytes derived from ASD individuals alter behavior and destabilize neuronal activity through aberrant Ca(2+) signaling. Mol Psychiatry 27, 2470–2484. 10.1038/s41380-022-01486-x.

21. Vakilzadeh, G., and Martinez-Cerdeno, V. (2023). Pathology and Astrocytes in Autism. Neuropsychiatr Dis Treat 19, 841–850. 10.2147/NDT.S390053.

22. Wang, Q., Kong, Y., Wu, D.Y., Liu, J.H., Jie, W., You, Q.L., Huang, L., Hu, J., Chu, H.D., Gao, F., et al. (2021). Impaired calcium signaling in astrocytes modulates autism spectrum disorder-like behaviors in mice. Nat Commun 12, 3321. 10.1038/s41467-021-23843-0.

23. Kahanovitch, U., Patterson, K.C., Hernandez, R., and Olsen, M.L. (2019). Glial Dysfunction in MeCP2 Deficiency Models: Implications for Rett Syndrome. Int J Mol Sci 20. 10.3390/ijms20153813.

24. Higashimori, H., Morel, L., Huth, J., Lindemann, L., Dulla, C., Taylor, A., Freeman, M., and Yang, Y. (2013). Astroglial FMRP-dependent translational down-regulation of mGluR5 underlies glutamate transporter GLT1 dysregulation in the fragile X mouse. Hum Mol Genet 22, 2041–2054. 10.1093/hmg/ddt055.

25. Higashimori, H., Schin, C.S., Chiang, M.S., Morel, L., Shoneye, T.A., Nelson, D.L., and Yang, Y. (2016). Selective Deletion of Astroglial FMRP Dysregulates Glutamate Transporter GLT1 and Contributes to Fragile X Syndrome Phenotypes In Vivo. J Neurosci 36, 7079–7094. 10.1523/JNEUROSCI.1069-16.2016.

26. Jacobs, S., and Doering, L.C. (2010). Astrocytes prevent abnormal neuronal development in the fragile x mouse. J Neurosci 30, 4508–4514. 10.1523/JNEUROSCI.5027-09.2010.

27. Jin, S.X., Higashimori, H., Schin, C., Tamashiro, A., Men, Y., Chiang, M.S.R., Jarvis, R., Cox, D., Feig, L., and Yang, Y. (2021). Astroglial FMRP modulates synaptic signaling and behavior phenotypes in FXS mouse model. Glia 69, 594–608. 10.1002/glia.23915.

28. Molofsky, A.V., Krencik, R., Ullian, E.M., Tsai, H.H., Deneen, B., Richardson, W.D., Barres, B.A., and Rowitch, D.H. (2012). Astrocytes and disease: a neurodevelopmental perspective. Genes Dev 26, 891–907. 10.1101/gad.188326.112.

29. Dong, Q., Kim, J., Nguyen, L., Bu, Q., and Chang, Q. (2020). An Astrocytic Influence on Impaired Tonic Inhibition in Hippocampal CA1 Pyramidal Neurons in a Mouse Model of Rett Syndrome. J Neurosci 40, 6250–6261. 10.1523/JNEUROSCI.3042-19.2020.

30. Garcia, O., Torres, M., Helguera, P., Coskun, P., and Busciglio, J. (2010). A role for thrombospondin-1 deficits in astrocyte-mediated spine and synaptic pathology in Down’s syndrome. PLoS One 5, e14200. 10.1371/journal.pone.0014200.

31. LeBlanc, J.J., DeGregorio, G., Centofante, E., Vogel-Farley, V.K., Barnes, K., Kaufmann, W.E., Fagiolini, M., and Nelson, C.A. (2015). Visual evoked potentials detect cortical processing deficits in Rett syndrome. Ann Neurol 78, 775–786. 10.1002/ana.24513.

32. Kostanian, D., Rebreikina, A., Voinova, V., and Sysoeva, O. (2023). Effect of presentation rate on auditory processing in Rett syndrome: event-related potential study. Mol Autism 14, 40. 10.1186/s13229-023-00566-1.

33. Foxe, J.J., Burke, K.M., Andrade, G.N., Djukic, A., Frey, H.P., and Molholm, S. (2016). Automatic cortical representation of auditory pitch changes in Rett syndrome. J Neurodev Disord 8, 34. 10.1186/s11689-016-9166-5.

34. Brima, T., Molholm, S., Molloy, C.J., Sysoeva, O.V., Nicholas, E., Djukic, A., Freedman, E.G., and Foxe, J.J. (2019). Auditory sensory memory span for duration is severely curtailed in females with Rett syndrome. Transl Psychiatry 9, 130. 10.1038/s41398-019-0463-0.

35. Roche, K.J., LeBlanc, J.J., Levin, A.R., O’Leary, H.M., Baczewski, L.M., and Nelson, C.A. (2019). Electroencephalographic spectral power as a marker of cortical function and disease severity in girls with Rett syndrome. J Neurodev Disord 11, 15. 10.1186/s11689-019-9275-z.

36. Saby, J.N., Peters, S.U., Roberts, T.P.L., Nelson, C.A., and Marsh, E.D. (2020). Evoked Potentials and EEG Analysis in Rett Syndrome and Related Developmental Encephalopathies: Towards a Biomarker for Translational Research. Front Integr Neurosci 14, 30. 10.3389/fnint.2020.00030.

37. Villemagne, P.M., Naidu, S., Villemagne, V.L., Yaster, M., Wagner, H.N., Jr., Harris, J.C., Moser, H.W., Johnston, M.V., Dannals, R.F., and Wong, D.F. (2002). Brain glucose metabolism in Rett Syndrome. Pediatr Neurol 27, 117–122. 10.1016/s0887-8994(02)00399-5.

38. Takeguchi, R., Kuroda, M., Tanaka, R., Suzuki, N., Akaba, Y., Tsujimura, K., Itoh, M., and Takahashi, S. (2022). Structural and functional changes in the brains of patients with Rett syndrome: A multimodal MRI study. J Neurol Sci 441, 120381. 10.1016/j.jns.2022.120381.

39. Yoshikawa, H., Kaga, M., Suzuki, H., Sakuragawa, N., and Arima, M. (1991). Giant somatosensory evoked potentials in the Rett syndrome. Brain Dev 13, 36–39. 10.1016/s0387-7604(12)80295-6.

40. Sysoeva, O., Maximenko, V., Kuc, A., Voinova, V., Martynova, O., and Hramov, A. (2023). Abnormal spectral and scale-free properties of resting-state EEG in girls with Rett syndrome. Sci Rep 13, 12932. 10.1038/s41598-023-39398-7.

41. Belichenko, P.V., Hagberg, B., and Dahlstrom, A. (1997). Morphological study of neocortical areas in Rett syndrome. Acta Neuropathol 93, 50–61. 10.1007/s004010050582.

42. Belichenko, P.V., Oldfors, A., Hagberg, B., and Dahlstrom, A. (1994). Rett syndrome: 3-D confocal microscopy of cortical pyramidal dendrites and afferents. Neuroreport 5, 1509–1513.

43. Dani, V.S., Chang, Q., Maffei, A., Turrigiano, G.G., Jaenisch, R., and Nelson, S.B. (2005). Reduced cortical activity due to a shift in the balance between excitation and inhibition in a mouse model of Rett syndrome. Proc Natl Acad Sci U S A 102, 12560–12565. 10.1073/pnas.0506071102.

44. Xu, X., Miller, E.C., and Pozzo-Miller, L. (2014). Dendritic spine dysgenesis in Rett syndrome. Front Neuroanat 8, 97. 10.3389/fnana.2014.00097.

45. Landi, S., Putignano, E., Boggio, E.M., Giustetto, M., Pizzorusso, T., and Ratto, G.M. (2011). The short-time structural plasticity of dendritic spines is altered in a model of Rett syndrome. Sci Rep 1, 45. 10.1038/srep00045.

46. Banerjee, A., Rikhye, R.V., Breton-Provencher, V., Tang, X., Li, C., Li, K., Runyan, C.A., Fu, Z., Jaenisch, R., and Sur, M. (2016). Jointly reduced inhibition and excitation underlies circuit-wide changes in cortical processing in Rett syndrome. Proc Natl Acad Sci U S A 113, E7287–E7296. 10.1073/pnas.1615330113.

47. Tropea, D., Giacometti, E., Wilson, N.R., Beard, C., McCurry, C., Fu, D.D., Flannery, R., Jaenisch, R., and Sur, M. (2009). Partial reversal of Rett Syndrome-like symptoms in MeCP2 mutant mice. Proc Natl Acad Sci U S A 106, 2029–2034. 10.1073/pnas.0812394106.

48. Fukuda, T., Itoh, M., Ichikawa, T., Washiyama, K., and Goto, Y. (2005). Delayed maturation of neuronal architecture and synaptogenesis in cerebral cortex of Mecp2-deficient mice. J Neuropathol Exp Neurol 64, 537–544. 10.1093/jnen/64.6.537.

49. Rietveld, L., Stuss, D.P., McPhee, D., and Delaney, K.R. (2015). Genotype-specific effects of Mecp2 loss-of-function on morphology of Layer V pyramidal neurons in heterozygous female Rett syndrome model mice. Front Cell Neurosci 9, 145. 10.3389/fncel.2015.00145.

50. Stuss, D.P., Boyd, J.D., Levin, D.B., and Delaney, K.R. (2012). MeCP2 mutation results in compartment-specific reductions in dendritic branching and spine density in layer 5 motor cortical neurons of YFP-H mice. PLoS One 7, e31896. 10.1371/journal.pone.0031896.

51. Kishi, N., and Macklis, J.D. (2004). MECP2 is progressively expressed in post-migratory neurons and is involved in neuronal maturation rather than cell fate decisions. Mol Cell Neurosci 27, 306–321. 10.1016/j.mcn.2004.07.006.

52. Guy, J., Gan, J., Selfridge, J., Cobb, S., and Bird, A. (2007). Reversal of neurological defects in a mouse model of Rett syndrome. Science 315, 1143–1147. 10.1126/science.1138389.

53. Garg, S.K., Lioy, D.T., Cheval, H., McGann, J.C., Bissonnette, J.M., Murtha, M.J., Foust, K.D., Kaspar, B.K., Bird, A., and Mandel, G. (2013). Systemic delivery of MeCP2 rescues behavioral and cellular deficits in female mouse models of Rett syndrome. J Neurosci 33, 13612–13620. 10.1523/JNEUROSCI.1854-13.2013.

54. Gadalla, K.K., Bailey, M.E., Spike, R.C., Ross, P.D., Woodard, K.T., Kalburgi, S.N., Bachaboina, L., Deng, J.V., West, A.E., Samulski, R.J., et al. (2013). Improved survival and reduced phenotypic severity following AAV9/MECP2 gene transfer to neonatal and juvenile male Mecp2 knockout mice. Mol Ther 21, 18–30. 10.1038/mt.2012.200.

55. Matagne, V., Ehinger, Y., Saidi, L., Borges-Correia, A., Barkats, M., Bartoli, M., Villard, L., and Roux, J.C. (2017). A codon-optimized Mecp2 transgene corrects breathing deficits and improves survival in a mouse model of Rett syndrome. Neurobiol Dis 99, 1–11. 10.1016/j.nbd.2016.12.009.

56. Matagne, V., Borloz, E., Ehinger, Y., Saidi, L., Villard, L., and Roux, J.C. (2021). Severe offtarget effects following intravenous delivery of AAV9-MECP2 in a female mouse model of Rett syndrome. Neurobiol Dis 149, 105235. 10.1016/j.nbd.2020.105235.

57. Na, E.S., Nelson, E.D., Adachi, M., Autry, A.E., Mahgoub, M.A., Kavalali, E.T., and Monteggia, L.M. (2012). A mouse model for MeCP2 duplication syndrome: MeCP2 overexpression impairs learning and memory and synaptic transmission. J Neurosci 32, 3109–3117. 10.1523/JNEUROSCI.6000-11.2012.

58. Lombardi, L.M., Baker, S.A., and Zoghbi, H.Y. (2015). MECP2 disorders: from the clinic to mice and back. J Clin Invest 125, 2914–2923. 10.1172/JCI78167.

59. Kolodkin, A.L., Matthes, D.J., and Goodman, C.S. (1993). The semaphorin genes encode a family of transmembrane and secreted growth cone guidance molecules. Cell 75, 1389–1399. 10.1016/0092-8674(93)90625-z.

60. Koropouli, E., and Kolodkin, A.L. (2014). Semaphorins and the dynamic regulation of synapse assembly, refinement, and function. Curr Opin Neurobiol 27, 1–7. 10.1016/j.conb.2014.02.005.

61. de Wit, J., and Verhaagen, J. (2003). Role of semaphorins in the adult nervous system. Prog Neurobiol 71, 249–267. 10.1016/j.pneurobio.2003.06.001.

62. Ding, J.B., Oh, W.J., Sabatini, B.L., and Gu, C. (2011). Semaphorin 3E-Plexin-D1 signaling controls pathway-specific synapse formation in the striatum. Nat Neurosci 15, 215–223. 10.1038/nn.3003.

63. Sahay, A., Kim, C.H., Sepkuty, J.P., Cho, E., Huganir, R.L., Ginty, D.D., and Kolodkin, A.L. (2005). Secreted semaphorins modulate synaptic transmission in the adult hippocampus. J Neurosci 25, 3613–3620. 10.1523/JNEUROSCI.5255-04.2005.

64. Tran, T.S., Rubio, M.E., Clem, R.L., Johnson, D., Case, L., Tessier-Lavigne, M., Huganir, R.L., Ginty, D.D., and Kolodkin, A.L. (2009). Secreted semaphorins control spine distribution and morphogenesis in the postnatal CNS. Nature 462, 1065–1069. 10.1038/nature08628.

65. Molofsky, A.V., Kelley, K.W., Tsai, H.H., Redmond, S.A., Chang, S.M., Madireddy, L., Chan, J.R., Baranzini, S.E., Ullian, E.M., and Rowitch, D.H. (2014). Astrocyte-encoded positional cues maintain sensorimotor circuit integrity. Nature 509, 189–194. 10.1038/nature13161.

66. Su, Y., Wang, X., Yang, Y., Chen, L., Xia, W., Hoi, K.K., Li, H., Wang, Q., Yu, G., Chen, X., et al. (2023). Astrocyte endfoot formation controls the termination of oligodendrocyte precursor cell perivascular migration during development. Neuron 111, 190–201 e198. 10.1016/j.neuron.2022.10.032.

67. Tamagnone, L., and Comoglio, P.M. (2000). Signalling by semaphorin receptors: cell guidance and beyond. Trends Cell Biol 10, 377–383. 10.1016/s0962-8924(00)01816-x.

68. Alto, L.T., and Terman, J.R. (2017). Semaphorins and their Signaling Mechanisms. Methods Mol Biol 1493, 1–25. 10.1007/978-1-4939-6448-2_1.

69. Renthal, W., Boxer, L.D., Hrvatin, S., Li, E., Silberfeld, A., Nagy, M.A., Griffith, E.C., Vierbuchen, T., and Greenberg, M.E. (2018). Characterization of human mosaic Rett syndrome brain tissue by single-nucleus RNA sequencing. Nat Neurosci 21, 1670–1679. 10.1038/s41593-018-0270-6.

70. Landucci, E., Brindisi, M., Bianciardi, L., Catania, L.M., Daga, S., Croci, S., Frullanti, E., Fallerini, C., Butini, S., Brogi, S., et al. (2018). iPSC-derived neurons profiling reveals GABAergic circuit disruption and acetylated alpha-tubulin defect which improves after iHDAC6 treatment in Rett syndrome. Exp Cell Res 368, 225–235. 10.1016/j.yexcr.2018.05.001.

71. Sanfeliu, A., Hokamp, K., Gill, M., and Tropea, D. (2019). Transcriptomic Analysis of Mecp2 Mutant Mice Reveals Differentially Expressed Genes and Altered Mechanisms in Both Blood and Brain. Front Psychiatry 10, 278. 10.3389/fpsyt.2019.00278.

72. Bajikar, S.S., Zhou, J., O’Hara, R., Tirumala, H.P., Durham, M.A., Trostle, A.J., Dias, M., Shao, Y., Chen, H., Wang, W., et al. (2025). Acute MeCP2 loss in adult mice reveals transcriptional and chromatin changes that precede neurological dysfunction and inform pathogenesis. Neuron 113, 380–395 e388. 10.1016/j.neuron.2024.11.006.

73. Matarazzo, V., and Ronnett, G.V. (2004). Temporal and regional differences in the olfactory proteome as a consequence of MeCP2 deficiency. Proc Natl Acad Sci U S A 101, 7763–7768. 10.1073/pnas.0307083101.

74. Sharifi, O., Haghani, V., Neier, K.E., Fraga, K.J., Korf, I., Hakam, S.M., Quon, G., Johansen, N., Yasui, D.H., and LaSalle, J.M. (2024). Sex-specific single cell-level transcriptomic signatures of Rett syndrome disease progression. Commun Biol 7, 1292. 10.1038/s42003-024-06990-0.

75. Bertoldi, M.L., Zalosnik, M.I., Fabio, M.C., Aja, S., Roth, G.A., Ronnett, G.V., and Degano, A.L. (2019). MeCP2 Deficiency Disrupts Kainate-Induced Presynaptic Plasticity in the Mossy Fiber Projections in the Hippocampus. Front Cell Neurosci 13, 286. 10.3389/fncel.2019.00286.

76. Degano, A.L., Pasterkamp, R.J., and Ronnett, G.V. (2009). MeCP2 deficiency disrupts axonal guidance, fasciculation, and targeting by altering Semaphorin 3F function. Mol Cell Neurosci 42, 243– 254. 10.1016/j.mcn.2009.07.009.

77. Belichenko, N.P., Belichenko, P.V., and Mobley, W.C. (2009). Evidence for both neuronal cell autonomous and nonautonomous effects of methyl-CpG-binding protein 2 in the cerebral cortex of female mice with Mecp2 mutation. Neurobiol Dis 34, 71–77. 10.1016/j.nbd.2008.12.016.

78. Samaco, R.C., McGraw, C.M., Ward, C.S., Sun, Y., Neul, J.L., and Zoghbi, H.Y. (2013). Female Mecp2(+/-) mice display robust behavioral deficits on two different genetic backgrounds providing a framework for pre-clinical studies. Hum Mol Genet 22, 96–109. 10.1093/hmg/dds406.

79. Krishnan, K., Wang, B.S., Lu, J., Wang, L., Maffei, A., Cang, J., and Huang, Z.J. (2015). MeCP2 regulates the timing of critical period plasticity that shapes functional connectivity in primary visual cortex. Proc Natl Acad Sci U S A 112, E4782–4791. 10.1073/pnas.1506499112.

80. Vogel Ciernia, A., Pride, M.C., Durbin-Johnson, B., Noronha, A., Chang, A., Yasui, D.H., Crawley, J.N., and LaSalle, J.M. (2017). Early motor phenotype detection in a female mouse model of Rett syndrome is improved by cross-fostering. Hum Mol Genet 26, 1839–1854. 10.1093/hmg/ddx087.

81. Wither, R.G., Lang, M., Zhang, L., and Eubanks, J.H. (2013). Regional MeCP2 expression levels in the female MeCP2-deficient mouse brain correlate with specific behavioral impairments. Exp Neurol 239, 49–59. 10.1016/j.expneurol.2012.09.005.

82. Mykins, M., Bridges, B., Jo, A., and Krishnan, K. (2024). Multidimensional Analysis of a Social Behavior Identifies Regression and Phenotypic Heterogeneity in a Female Mouse Model for Rett Syndrome. J Neurosci 44. 10.1523/JNEUROSCI.1078-23.2023.

83. Dong, H.W., Erickson, K., Lee, J.R., Merritt, J., Fu, C., and Neul, J.L. (2020). Detection of neurophysiological features in female R255X MeCP2 mutation mice. Neurobiol Dis 145, 105083. 10.1016/j.nbd.2020.105083.

84. Lau, B.Y.B., Krishnan, K., Huang, Z.J., and Shea, S.D. (2020). Maternal Experience-Dependent Cortical Plasticity in Mice Is Circuit- and Stimulus-Specific and Requires MECP2. J Neurosci 40, 1514–1526. 10.1523/JNEUROSCI.1964-19.2019.

85. Ribeiro, M.C., and MacDonald, J.L. (2020). Sex differences in Mecp2-mutant Rett syndrome model mice and the impact of cellular mosaicism in phenotype development. Brain Res 1729, 146644. 10.1016/j.brainres.2019.146644.

86. Sharifi, O., and Yasui, D.H. (2021). The Molecular Functions of MeCP2 in Rett Syndrome Pathology. Front Genet 12, 624290. 10.3389/fgene.2021.624290.

87. Ruediger, T., Zimmer, G., Barchmann, S., Castellani, V., Bagnard, D., and Bolz, J. (2013). Integration of opposing semaphorin guidance cues in cortical axons. Cereb Cortex 23, 604–614. 10.1093/cercor/bhs044.

88. Polleux, F., Morrow, T., and Ghosh, A. (2000). Semaphorin 3A is a chemoattractant for cortical apical dendrites. Nature 404, 567–573. 10.1038/35007001.

89. Niquille, M., Garel, S., Mann, F., Hornung, J.P., Otsmane, B., Chevalley, S., Parras, C., Guillemot, F., Gaspar, P., Yanagawa, Y., and Lebrand, C. (2009). Transient neuronal populations are required to guide callosal axons: a role for semaphorin 3C. PLoS Biol 7, e1000230. 10.1371/journal.pbio.1000230.

90. Martins, L.F., Brambilla, I., Motta, A., de Pretis, S., Bhat, G.P., Badaloni, A., Malpighi, C., Amin, N.D., Imai, F., Almeida, R.D., et al. (2022). Motor neurons use push-pull signals to direct vascular remodeling critical for their connectivity. Neuron 110, 4090–4107 e4011. 10.1016/j.neuron.2022.09.021.

91. Yang, W.J., Hu, J., Uemura, A., Tetzlaff, F., Augustin, H.G., and Fischer, A. (2015). Semaphorin-3C signals through Neuropilin-1 and PlexinD1 receptors to inhibit pathological angiogenesis. EMBO Mol Med 7, 1267–1284. 10.15252/emmm.201404922.

92. Guy, J., Hendrich, B., Holmes, M., Martin, J.E., and Bird, A. (2001). A mouse Mecp2-null mutation causes neurological symptoms that mimic Rett syndrome. Nat Genet 27, 322–326. 10.1038/85899.

93. Kishi, N., and Macklis, J.D. (2010). MeCP2 functions largely cell-autonomously, but also non-cell-autonomously, in neuronal maturation and dendritic arborization of cortical pyramidal neurons. Exp Neurol 222, 51–58. 10.1016/j.expneurol.2009.12.007.

94. Jentarra, G.M., Olfers, S.L., Rice, S.G., Srivastava, N., Homanics, G.E., Blue, M., Naidu, S., and Narayanan, V. (2010). Abnormalities of cell packing density and dendritic complexity in the MeCP2 A140V mouse model of Rett syndrome/X-linked mental retardation. BMC Neurosci 11, 19. 10.1186/1471-2202-11-19.

95. Durand, S., Patrizi, A., Quast, K.B., Hachigian, L., Pavlyuk, R., Saxena, A., Carninci, P., Hensch, T.K., and Fagiolini, M. (2012). NMDA receptor regulation prevents regression of visual cortical function in the absence of Mecp2. Neuron 76, 1078–1090. 10.1016/j.neuron.2012.12.004.

96. Plein, A., Calmont, A., Fantin, A., Denti, L., Anderson, N.A., Scambler, P.J., and Ruhrberg, C. (2015). Neural crest-derived SEMA3C activates endothelial NRP1 for cardiac outflow tract septation. J Clin Invest 125, 2661–2676. 10.1172/JCI79668.

97. Carter, J.C., Lanham, D.C., Pham, D., Bibat, G., Naidu, S., and Kaufmann, W.E. (2008). Selective cerebral volume reduction in Rett syndrome: a multiple-approach MR imaging study. AJNR Am J Neuroradiol 29, 436–441. 10.3174/ajnr.A0857.

98. Jan, T.Y., Wong, L.C., Hsu, C.J., Huang, C.J., Peng, S.S., Tseng, W.I., and Lee, W.T. (2024). Developmental change of brain volume in Rett syndrome in Taiwan. J Neurodev Disord 16, 36. 10.1186/s11689-024-09549-6.

99. Allemang-Grand, R., Ellegood, J., Spencer Noakes, L., Ruston, J., Justice, M., Nieman, B.J., and Lerch, J.P. (2017). Neuroanatomy in mouse models of Rett syndrome is related to the severity of Mecp2 mutation and behavioral phenotypes. Mol Autism 8, 32. 10.1186/s13229-017-0138-8.

100. Bittolo, T., Raminelli, C.A., Deiana, C., Baj, G., Vaghi, V., Ferrazzo, S., Bernareggi, A., and Tongiorgi, E. (2016). Pharmacological treatment with mirtazapine rescues cortical atrophy and respiratory deficits in MeCP2 null mice. Sci Rep 6, 19796. 10.1038/srep19796.

101. Wood, L., and Shepherd, G.M. (2010). Synaptic circuit abnormalities of motor-frontal layer 2/3 pyramidal neurons in a mutant mouse model of Rett syndrome. Neurobiol Dis 38, 281–287. 10.1016/j.nbd.2010.01.018.

102. Belichenko, N.P., Belichenko, P.V., Li, H.H., Mobley, W.C., and Francke, U. (2008). Comparative study of brain morphology in Mecp2 mutant mouse models of Rett syndrome. J Comp Neurol 508, 184–195. 10.1002/cne.21673.

103. Metcalf, B.M., Mullaney, B.C., Johnston, M.V., and Blue, M.E. (2006). Temporal shift in methyl-CpG binding protein 2 expression in a mouse model of Rett syndrome. Neuroscience 139, 1449–1460. 10.1016/j.neuroscience.2006.01.060.

104. Chapleau, C.A., Calfa, G.D., Lane, M.C., Albertson, A.J., Larimore, J.L., Kudo, S., Armstrong, D.L., Percy, A.K., and Pozzo-Miller, L. (2009). Dendritic spine pathologies in hippocampal pyramidal neurons from Rett syndrome brain and after expression of Rett-associated MECP2 mutations. Neurobiol Dis 35, 219–233. 10.1016/j.nbd.2009.05.001.

105. Risher, W.C., Ustunkaya, T., Singh Alvarado, J., and Eroglu, C. (2014). Rapid Golgi analysis method for efficient and unbiased classification of dendritic spines. PLoS One 9, e107591. 10.1371/journal.pone.0107591.

106. Mierau, S.B., Patrizi, A., Hensch, T.K., and Fagiolini, M. (2016). Cell-Specific Regulation of N-Methyl-D-Aspartate Receptor Maturation by Mecp2 in Cortical Circuits. Biol Psychiatry 79, 746–754. 10.1016/j.biopsych.2015.05.018.

107. Santos, M., Silva-Fernandes, A., Oliveira, P., Sousa, N., and Maciel, P. (2007). Evidence for abnormal early development in a mouse model of Rett syndrome. Genes Brain Behav 6, 277–286. 10.1111/j.1601-183X.2006.00258.x.

108. Segawa, M. (2005). Early motor disturbances in Rett syndrome and its pathophysiological importance. Brain Dev 27 *Suppl 1*, S54–S58. 10.1016/j.braindev.2004.11.010.

109. von Tetzchner, S., Jacobsen, K.H., Smith, L., Skjeldal, O.H., Heiberg, A., and Fagan, J.F. (1996). Vision, cognition and developmental characteristics of girls and women with Rett syndrome. Dev Med Child Neurol 38, 212–225. 10.1111/j.1469-8749.1996.tb15083.x.

110. Prusky, G.T., Alam, N.M., Beekman, S., and Douglas, R.M. (2004). Rapid quantification of adult and developing mouse spatial vision using a virtual optomotor system. Invest Ophthalmol Vis Sci 45, 4611–4616. 10.1167/iovs.04-0541.

111. Crawley, J.N. (2008). Behavioral phenotyping strategies for mutant mice. Neuron 57, 809–818. 10.1016/j.neuron.2008.03.001.

112. Christie, S.M., Hao, J., Tracy, E., Buck, M., Yu, J.S., and Smith, A.W. (2021). Interactions between semaphorins and plexin-neuropilin receptor complexes in the membranes of live cells. J Biol Chem 297, 100965. 10.1016/j.jbc.2021.100965.

113. Chen, H., He, Z., Bagri, A., and Tessier-Lavigne, M. (1998). Semaphorin-neuropilin interactions underlying sympathetic axon responses to class III semaphorins. Neuron 21, 1283–1290. 10.1016/s0896-6273(00)80648-0.

114. Gitler, A.D., Lu, M.M., and Epstein, J.A. (2004). PlexinD1 and semaphorin signaling are required in endothelial cells for cardiovascular development. Dev Cell 7, 107–116. 10.1016/j.devcel.2004.06.002.

115. Cheng, S., Butrus, S., Tan, L., Xu, R., Sagireddy, S., Trachtenberg, J.T., Shekhar, K., and Zipursky, S.L. (2022). Vision-dependent specification of cell types and function in the developing cortex. Cell 185, 311–327 e324. 10.1016/j.cell.2021.12.022.

116. Ferrario, J.E., Baskaran, P., Clark, C., Hendry, A., Lerner, O., Hintze, M., Allen, J., Chilton, J.K., and Guthrie, S. (2012). Axon guidance in the developing ocular motor system and Duane retraction syndrome depends on Semaphorin signaling via alpha2-chimaerin. Proc Natl Acad Sci U S A 109, 14669–14674. 10.1073/pnas.1116481109.

117. Brown, C.B., Feiner, L., Lu, M.M., Li, J., Ma, X., Webber, A.L., Jia, L., Raper, J.A., and Epstein, J.A. (2001). PlexinA2 and semaphorin signaling during cardiac neural crest development. Development 128, 3071–3080. 10.1242/dev.128.16.3071.

118. Quach, T.T., Stratton, H.J., Khanna, R., Kolattukudy, P.E., Honnorat, J., Meyer, K., and Duchemin, A.M. (2021). Intellectual disability: dendritic anomalies and emerging genetic perspectives. Acta Neuropathol 141, 139–158. 10.1007/s00401-020-02244-5.

119. Wang, Q., Chiu, S.L., Koropouli, E., Hong, I., Mitchell, S., Easwaran, T.P., Hamilton, N.R., Gustina, A.S., Zhu, Q., Ginty, D.D., et al. (2017). Neuropilin-2/PlexinA3 Receptors Associate with GluA1 and Mediate Sema3F-Dependent Homeostatic Scaling in Cortical Neurons. Neuron 96, 1084–1098 e1087. 10.1016/j.neuron.2017.10.029.

120. McDermott, J.E., Goldblatt, D., and Paradis, S. (2018). Class 4 Semaphorins and Plexin-B receptors regulate GABAergic and glutamatergic synapse development in the mammalian hippocampus. Mol Cell Neurosci 92, 50–66. 10.1016/j.mcn.2018.06.008.

121. Acker, D.W.M., Wong, I., Kang, M., and Paradis, S. (2018). Semaphorin 4D promotes inhibitory synapse formation and suppresses seizures in vivo. Epilepsia 59, 1257–1268. 10.1111/epi.14429.

122. Kuzirian, M.S., Moore, A.R., Staudenmaier, E.K., Friedel, R.H., and Paradis, S. (2013). The class 4 semaphorin Sema4D promotes the rapid assembly of GABAergic synapses in rodent hippocampus. J Neurosci 33, 8961–8973. 10.1523/JNEUROSCI.0989-13.2013.

123. Ure, K., Lu, H., Wang, W., Ito-Ishida, A., Wu, Z., He, L.J., Sztainberg, Y., Chen, W., Tang, J., and Zoghbi, H.Y. (2016). Restoration of Mecp2 expression in GABAergic neurons is sufficient to rescue multiple disease features in a mouse model of Rett syndrome. Elife 5. 10.7554/eLife.14198.

124. Rohm, B., Ottemeyer, A., Lohrum, M., and Puschel, A.W. (2000). Plexin/neuropilin complexes mediate repulsion by the axonal guidance signal semaphorin 3A. Mech Dev 93, 95–104. 10.1016/s0925-4773(00)00269-0.

125. Gay, C.M., Zygmunt, T., and Torres-Vazquez, J. (2011). Diverse functions for the semaphorin receptor PlexinD1 in development and disease. Dev Biol 349, 1–19. 10.1016/j.ydbio.2010.09.008.

126. Ito, Y., Oinuma, I., Katoh, H., Kaibuchi, K., and Negishi, M. (2006). Sema4D/plexin-B1 activates GSK-3beta through R-Ras GAP activity, inducing growth cone collapse. EMBO Rep 7, 704–709. 10.1038/sj.embor.7400737.

127. Oinuma, I., Ishikawa, Y., Katoh, H., and Negishi, M. (2004). The Semaphorin 4D receptor Plexin-B1 is a GTPase activating protein for R-Ras. Science 305, 862–865. 10.1126/science.1097545.

128. Oinuma, I., Katoh, H., and Negishi, M. (2006). Semaphorin 4D/Plexin-B1-mediated R-Ras GAP activity inhibits cell migration by regulating beta(1) integrin activity. J Cell Biol 173, 601–613. 10.1083/jcb.200508204.

129. Oinuma, I., Katoh, H., and Negishi, M. (2004). Molecular dissection of the semaphorin 4D receptor plexin-B1-stimulated R-Ras GTPase-activating protein activity and neurite remodeling in hippocampal neurons. J Neurosci 24, 11473–11480. 10.1523/JNEUROSCI.3257-04.2004.

130. Uesugi, K., Oinuma, I., Katoh, H., and Negishi, M. (2009). Different requirement for Rnd GTPases of R-Ras GAP activity of Plexin-C1 and Plexin-D1. J Biol Chem 284, 6743–6751. 10.1074/jbc.M805213200.

131. Assous, M., Martinez, E., Eisenberg, C., Shah, F., Kosc, A., Varghese, K., Espinoza, D., Bhimani, S., Tepper, J.M., Shiflett, M.W., and Tran, T.S. (2019). Neuropilin 2 Signaling Mediates Corticostriatal Transmission, Spine Maintenance, and Goal-Directed Learning in Mice. J Neurosci 39, 8845–8859. 10.1523/JNEUROSCI.1006-19.2019.

132. Shiflett, M.W., Gavin, M., and Tran, T.S. (2015). Altered hippocampal-dependent memory and motor function in neuropilin 2-deficient mice. Transl Psychiatry 5, e521. 10.1038/tp.2015.17.

133. Gu, C., Yoshida, Y., Livet, J., Reimert, D.V., Mann, F., Merte, J., Henderson, C.E., Jessell, T.M., Kolodkin, A.L., and Ginty, D.D. (2005). Semaphorin 3E and plexin-D1 control vascular pattern independently of neuropilins. Science 307, 265–268. 10.1126/science.1105416.

134. Chauvet, S., Cohen, S., Yoshida, Y., Fekrane, L., Livet, J., Gayet, O., Segu, L., Buhot, M.C., Jessell, T.M., Henderson, C.E., and Mann, F. (2007). Gating of Sema3E/PlexinD1 signaling by neuropilin-1 switches axonal repulsion to attraction during brain development. Neuron 56, 807–822. 10.1016/j.neuron.2007.10.019.

135. De Rubeis, S., He, X., Goldberg, A.P., Poultney, C.S., Samocha, K., Cicek, A.E., Kou, Y., Liu, L., Fromer, M., Walker, S., et al. (2014). Synaptic, transcriptional and chromatin genes disrupted in autism. Nature 515, 209–215. 10.1038/nature13772.

136. Stessman, H.A., Xiong, B., Coe, B.P., Wang, T., Hoekzema, K., Fenckova, M., Kvarnung, M., Gerdts, J., Trinh, S., Cosemans, N., et al. (2017). Targeted sequencing identifies 91 neurodevelopmental-disorder risk genes with autism and developmental-disability biases. Nat Genet 49, 515–526. 10.1038/ng.3792.

137. Menon, L., and Mihailescu, M.R. (2007). Interactions of the G quartet forming semaphorin 3F RNA with the RGG box domain of the fragile X protein family. Nucleic Acids Res 35, 5379–5392. 10.1093/nar/gkm581.

